# A hierarchy of migratory keratinocytes maintains the tympanic membrane

**DOI:** 10.1101/687947

**Authors:** Stacey M. Frumm, Kevin Shengyang Yu, Joseph Chang, Jordan A. Briscoe, Katharine P. Lee, Lauren E. Byrnes, Julie B. Sneddon, Aaron D. Tward

## Abstract

Although the conductive function of the tympanic membrane (TM) is critical for hearing, it is unknown how the organ maintains cellular homeostasis. Using a combination of single-cell RNA sequencing, lineage tracing, whole-organ explant, and live-cell imaging, we demonstrate that the stem cells of the TM epidermis reside in a distinct location at the superior portion of the TM and, as progeny migrate inferiorly, Pdgfra+ fibroblasts maintain a niche supporting proliferation of committed progenitors, while keratinocytes distal from the niche differentiate. Thus, the TM has a three dimensional differentiation hierarchy of keratinocytes distinct from that at other epidermal sites. The TM represents a physiological context where, in the absence of injury, keratinocytes both transit through a proliferative committed progenitor state and exhibit directional lateral migration. This work forms a foundation for understanding common disorders of the TM and introduces a new model system for the understanding of keratinocyte biology.

## Introduction

Disorders of the tympanic membrane (TM), also known as the eardrum, and middle ear space are common and morbid. There are over 750,000 surgical procedures performed on the TM each year (Hall et al. 2017), and in 2006 there were 8,000,000 physicians visits by children for ear infections (Soni 2008). Disorders of the TM including perforation, retraction, cholesteatoma, tympanosclerosis, bullous myringitis, and keratosis obturans can lead to decreased hearing, chronic infections and drainage from the ear, vertigo, and, in severe cases, meningitis or death. Despite its medical importance, little is known about the cellular composition of the TM, or how cellular homeostasis of this important organ is maintained.

The TM is a central component of the conductive apparatus of the ear (**Figure 1A**). It transmits sound vibrations from the external auditory canal (EAC) to the middle ear ossicles, which in turn transfer those vibrations to the cochlea. The TM is comprised of two distinct anatomic regions (**Figure 1B**). The pars tensa is the inferior portion and vibrates in response to sound waves. The handle of the first middle ear ossicle, the malleus, is embedded within the mesenchymal layer of the pars tensa (**Figure 1C**). The pars flaccida is the superior portion of the TM and is believed to contribute less to sound transmission. The outer layer of the TM is epidermis continuous with that of the EAC. The TM is a somewhat unusual site of epidermis, as it does not possess adnexal structures such as hair follicles or sweat glands. Thus, it provides a unique site to study epidermal cellular dynamics in the absence of confounding from other skin stem cell populations.

**Figure 1:**
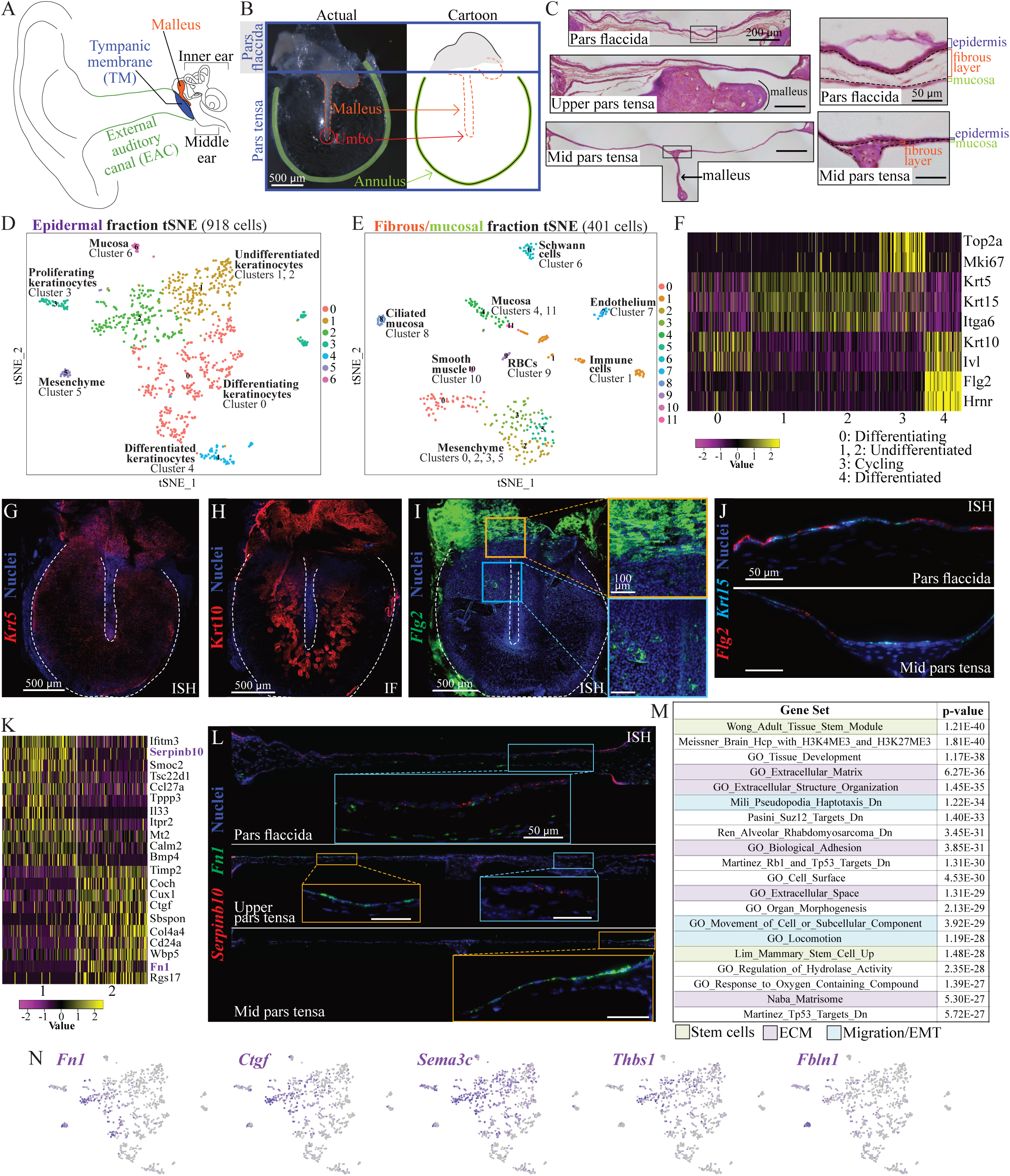
Single-cell RNA sequencing (scRNA-Seq) identifies cell types in the murine TM. (A) Diagram of the ear with key structures annotated. The tympanic membrane (TM) is located between the external auditory canal (EAC) and middle ear. (B) Image of a murine TM dissected en bloc and a cartoon representation with key anatomy annotated. (C) Sections of the TM at the level of the pars flaccida (top), upper pars tensa (middle) and mid pars tensa (bottom). The epidermal layer is oriented upward in all. The indicated areas of the pars flaccida and mid pars tensa are shown magnified to the right, with the tissue layers (epidermis, fibrous layer, and mucosa) indicated. In all three sections on the left, the scale bar indicates 200 μm, and in both sections on the right, the scale bar indicates 50 μm. (D) t-Distributed Stochastic Neighbor Embedding (tSNE) visualization of the cell clusters identified in the epidermal fraction. (E) tSNE visualization of the cell clusters identified in the fibrous/mucosal fraction. (F) Heat-map showing expression of genes associated with keratinocyte differentiation states in keratinocyte clusters 0 - 4. Yellow indicates high expression and violet low expression. Each column is a single cell. (G) *In situ* hybridization (ISH) showing expression of *Krt5* (red) in a whole-mount TM. (H) Immunofluorescence (IF) for Krt10 (red) in a whole-mount TM. (I) ISH showing expression of *Flg2* (green) in a whole-mount TM. Yellow and blue boxes indicate areas shown at higher magnification on the right. The yellow box highlights an area at the pars flaccida/pars tensa junction, and the blue box highlights an area of the pars tensa over the malleus. The scale bars in both magnified images on the right indicate 100 μm. (J) ISH showing expression of *Flg2* (red) and *Krt15* (cyan) in TM sections from the pars flaccida (top) and pars tensa (bottom). The epidermal layer is oriented upward. Both scale bars indicate 50 μm. (K) Heat-map showing genes differentially expressed between undifferentiated keratinocyte clusters 1 and 2. (L) ISH showing expression of *Serpinb10* (red) and *Fn1* (green) in TM sections from the pars flaccida (top, ∼1500 μm across), upper pars tensa (middle, ∼1650 μm across), and mid pars tensa (bottom, ∼1800 μm across). Blue and yellow boxes highlight areas shown at greater magnification as insets; the scale bars in all insets indicate 50 μm. The epidermal layer is oriented upward in all images. (M) Top gene sets from MSigDB that overlap with the 100 genes most upregulated in cluster 2 relative to cluster 1. (N) tSNE plots depicting expression of *Fn1, Ctgf, Sema3c, Thbs1*, and *Fbln1* in each cell from the epidermal fraction. Grey dots indicate cells in which no transcript was detected.

In addition to performing the barrier function of any epidermis, the epidermis of the TM and EAC must also prevent the accumulation of cellular and exogenous debris via a cleaning mechanism. Since the EAC terminates in a blind sac at the TM, if differentiated superficial keratinocytes were simply sloughed into the EAC, then the EAC would fill up with cellular debris. This would prevent sound transmission, create a fertile environment for infection, and cause erosion of surrounding critical structures. Indeed, variations of this situation occur in patients with keratosis obturans, cholesteatoma, and cerumen impaction (Soucek & Michaels 1993; Corbridge et al. 1996; Kuo et al. 2015; Marchisio et al. 2016). Previous experiments have demonstrated that ink spots placed on the TMs of mammals, including humans, migrate laterally along the TM and down the EAC (Stinson 1936; Litton 1963; Alberti 1964; O’Donoghue 1984; Kakoi et al. 1996; Tinling & Chole 2006). Pulse-chase experiments with the thymidine analog BrdU have indicated that proliferation of cells in the TM is not uniform, but rather restricted to a few locations, and that newly proliferated cells likely exhibit migratory behavior (Kakoi & Anniko 1997; Koba 1995). Additional studies have demonstrated clonogenic capacity and migratory behavior of epithelial cells derived from the mouse and human TM *in vitro* (Redmond et al. 2011; Liew et al. 2017; Ong et al. 2017; Liew et al. 2018). It is further clear that following perforation of the TM there is a massive cellular response typically leading to the rapid healing in all mammals, including humans, investigated thus far (Santa Maria et al. 2010; Rollin et al. 2011; Lou et al. 2012; Chari et al. 2019). However, our knowledge of the cellular identities, dynamics, and mechanisms of TM homeostasis is otherwise limited.

At other anatomical sites, the epidermis is maintained by a population of basal keratinocyte stem cells (SCs) that are capable of both self-renewal as well as differentiation, and that exhibit neutral competition kinetics in most homeostatic contexts studied to date (Clayton et al. 2007; Doupé et al. 2010; Mascré et al. 2012; Lim et al. 2013). In addition, some evidence exists for a hierarchical differentiation scheme in uninjured skin whereby the SCs give rise to a proliferative committed progenitor (CP) or transit amplifying population that has a decreased propensity for long term maintenance of the tissue (Mascré et al. 2012; Jones & Simons 2008; Sada et al. 2016). Part of the challenge in distinguishing between these models in epidermis is that both the SC and putative CP populations exist in similar locations and express similar markers, and thus the strongest evidence for the existence of keratinocyte CPs comes from mathematical modeling of the fates of clonal progeny (Mascré et al. 2012). Additionally, the daughter cells of proliferating skin keratinocytes either remain in the basal layer or commit to differentiation and progressively move to more apical layers of the epidermis. Although basal patches of SC progeny arise from individual marked SCs over time, it is believed that, in uninjured skin, basal keratinocytes do not exhibit active lateral migratory behavior in the plane of the basement membrane (Rompolas et al. 2016). However, after wounding, basal keratinocytes (and likely suprabasal keratinocytes) undergo a transcriptional shift and acquire the property of active and directional lateral migration (Gurtner et al. 2008; Park et al. 2017; Grinnell 1990). The TM epidermis is thus distinct, as the ink dot studies suggest that lateral migration occurs in homeostasis as well as injury. How this impacts tissue maintenance, and whether progenitor populations equivalent to those seen at other epidermal sites exist in the TM, has not previously been evaluated.

In this work, we combine single-cell RNA sequencing, live cell imaging, lineage tracing, and a novel explant model to explore the biology of the TM. We demonstrate that, under homeostasis, the TM possesses at least three distinct populations of basal keratinocytes: a SC population arising from the junction between the pars tensa and pars flaccida with long-term renewal capability, a CP population near the handle of the malleus in the pars tensa that exhibits proliferative and migratory capacity but lacks long-term renewal potential, and a migratory population throughout the majority of the pars tensa that does not proliferate. Furthermore, we describe a system to maintain whole murine TMs in culture, and identify Pdgf signaling as required for maintaining regional proliferation of TM keratinocytes.

## Results

### Identifying cell populations in the murine TM

Prior work has described three basic TM layers—the epidermis, fibrous layer, and mucosa—but not the full diversity of TM cell types (**Figure 1C**). We sought to define the resident cell populations of the TM via single-cell RNA sequencing (scRNA-Seq) using the 10x Genomics Single Cell Solution v2 platform. Thirty-eight adult murine TMs were pooled and split into two anatomic fractions, epidermal and fibrous/mucosal, to optimally dissociate different cell types. Two corresponding datasets were generated and analyzed with Seurat (**Figure 1D-E**) (Satija et al. 2015).

In the epidermal and fibrous/mucosal fractions, respectively, 918 and 401 cells were captured (**Figure 1D-E**). In the epidermal fraction, seven cell clusters were identified, including five keratinocyte clusters (numbered 0, 1, 2, 3, and 4), as well as mesenchymal (5) and mucosal (6) cells (**Figure 1D**). Keratinocytes in cluster 3 are cycling, marked by expression of *Top2a* and *Mki67* (**Figure 1F**); almost no cycling cells are seen in the other tissue layers (**Figure S1**). The prominent axis of distinction among the other keratinocyte clusters is differentiation state. Clusters 1 and 2 are relatively undifferentiated; they express markers of basal keratinocytes, including *Krt5, Krt15*, and *Itga6* (**Figure 1F**). Cluster 0 is an intermediate differentiation state; the cells have lost expression of the basal-like markers and gained expression of differentiation-associated genes, including *Krt10* and *Ivl*. Lastly, cluster 4 is terminally differentiated keratinocytes, expressing markers including *Flg2* and *Hrnr*. To determine the locations of these populations, both *in situ* hybridization (ISH) and immunofluorescence were utilized. *Krt5*+ cells are seen diffusely throughout the TM, while Krt10+ cells are predominantly located in the intermediate region of the pars tensa—although not over the malleus—as well as the pars flaccida (**Figure 1G-H**). *Flg2* is strongly expressed in the pars flaccida, while only occasional cells in the pars tensa express this gene (**Figure 1I**). In a section through the pars flaccida, robust *Flg2* expression is again seen in many cells, often overlying less differentiated *Krt15*+ keratinocytes. In a section through the pars tensa at a level where little stratification is seen, a patch of several *Flg2*+ cells is observed in the one-cell-thick epidermis (**Figure 1J**). Thus, the pars flaccida exhibits both differentiation and stratification, whereas the pars tensa exhibits minimal stratification, but preserves aspects of predominantly early differentiation. Overall, the gene expression profiles obtained among the keratinocytes were similar to those previously observed in other interfollicular epidermal sites (Cheng et al. 2018; Joost et al. 2016; Joost et al. 2018; Aragona et al. 2017).

Next, we identified genes whose expression distinguished between clusters 1 and 2—both relatively undifferentiated keratinocyte populations—and used these markers to visualize the two cell types in TM tissue. Of the genes differentially expressed, *Serpinb10* and *Fn1* were identified as discriminating markers of these two groups (**Figure 1K**). ISH to detect these transcripts in TM sections confirmed that their expression is not overlapping (**Figure 1L**). Keratinocytes in the pars flaccida predominantly express *Serpinb10*; *Fn1* is expressed in this region of the tissue, but by fibroblasts. In the pars tensa, a section through the middle of the malleus has only an *Fn1*+ keratinocyte population near the annulus, while a section through the upper pars tensa has separate *Fn1*+ and *Serpinb10*+ regions. Mixed regions of adjacent *Serpinb10*+ and *Fn1*+ keratinocytes were not observed. *Fn1* expression was looked at more broadly in whole-mount TMs (**Figure S2**). Regions of *Fn1*+ cells are seen near the annulus and over the malleus. Thus, clusters 1 and 2 mark distinct populations of basal keratinocytes with one found predominantly in the pars flaccida (*Serpinb10*+) and the other found predominantly in the pars tensa (*Fn1*+).

To further understand the functional distinctions between cells of clusters 1 and 2, we looked for gene sets in the Molecular Signatures Database that significantly overlapped with the genes most differentially expressed between the two clusters (Subramanian et al. 2005; Liberzon et al. 2011; Liberzon et al. 2015). For the genes more highly expressed in cluster 1, the most common terms enriched were related to ribosome function and biogenesis (**Table S1**). For the genes more highly expressed in cluster 2, many of the enriched terms were related to cell migration and the extracellular matrix (**Figure 1M** and **Table S2**). Among the genes more highly expressed in cluster 2 were *Fn1, Ctgf* (Kiwanuka et al. 2013), *Sema3c* (Curreli et al. 2016), *Thbs1* (Borsotti et al. 2015), and *Fbln1* (Feitosa et al. 2012), which are characteristic of the migratory cell populations (**Figure 1N**).

The same approaches were utilized to define and locate the cell populations in the fibrous/mucosal fraction, where 12 clusters were discovered. Among these are multiple *Vimentin*+ (*Vim*) mesenchymal populations, including: endothelium (*Pecam1*+), smooth muscle (*Acta2*+), and Schwann cells (*Mbp*+) (**Figure S3**). Four additional clusters of mesenchymal cells were identified, numbered 0, 2, 3, and 5. Inspection of the differentially expressed genes revealed that cluster 0 is *Gpx3*+, and clusters 2, 3, and 5 are *Igfbp3*+ (**Figure S4A-B**). ISH confirmed that *Gpx3* and *Igfbp3* do indeed mark distinct populations of fibroblasts (**Figure S4C**). The *Gpx3*+ cells appear to reside in proximity to bony elements—specifically, in the annular region and near the malleus. The *Igfbp3*+ cells are more diffusely spread throughout the fibrous layer of the TM.

Lastly, *Sox2* is a known marker of mucosa, and is expressed in both non-ciliated (clusters 4 and 11) and ciliated (cluster 8) mucosal cell clusters (**Figure S5A**)(Tucker et al. 2018). Cluster 8 is identifiable as ciliated mucosa by expression of motor proteins, including *Dnah5* and *Dynlrb2* (**Figure S5B**). The mucosal clusters express *Krt19*, while the keratinocyte clusters do not, and restriction of this keratin to the mucosal cells was confirmed by lineage tracing (**Figure S5C**). Of note, Krt19 has been proposed as a marker of TM epidermal stem cells in human and rat (Knutsson et al. 2011; Kim et al. 2015).

### scRNA-Seq of human TM tissue

Having cataloged the cell populations in the murine TM, we next addressed whether equivalent cell populations are also present in the human TM. Freshly isolated normal TM tissue obtained during a surgical approach for a skull base tumor was mechanically separated into more lateral and medial portions, then dissociated and subjected to scRNA-Seq. From the lateral portion, 1,157 cells were captured and 14 clusters identified, and from the medial portion, 2,708 cells were captured and 22 clusters identified (**Figure 2A**). Similar cell types were identified in the human dataset as in the murine dataset, confirming that the mouse is a valid model organism for study. These cell populations included keratinocytes (*KRT14*+), mesenchyme (*FBLN1*+), mucosa (*SOX2*+), and smooth muscle (*MYH11*+) (**Figure 2B**). Only approximately 200 keratinocytes were captured in each portion, but evidence of multiple differentiation states is exemplified by differential expression of *KRT15* and *KRT1* (**Figure 2C**).

**Figure 2:**
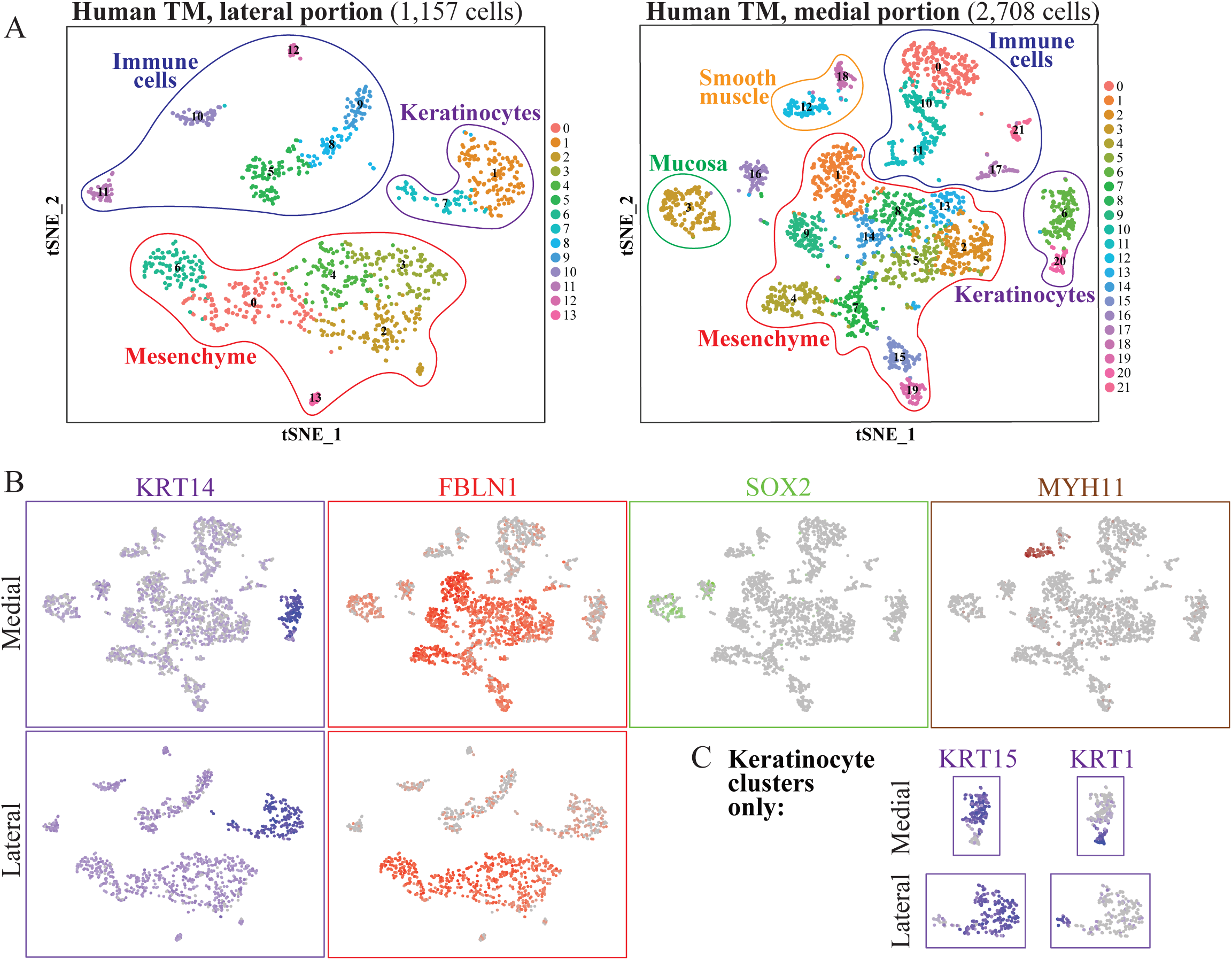
scRNA-Seq of the human TM. (A) tSNE plots of the cell clusters identified in scRNA-Seq of a human TM. The TM tissue was divided into more lateral and medial portions prior to dissociation, and the corresponding tSNE plots are on the left and right, respectively. (B) tSNE plots showing expression of *KRT14* (purple; keratinocytes), *FBLN1* (red; fibroblasts), *SOX2* (green; mucosa), and *MYH11* (brown; smooth muscle) in the two datasets. (C) Expression of *KRT15* and *KRT1* in the keratinocyte clusters.

### EdU incorporation demonstrates rapid turnover of the TM epidermis

We next wanted to characterize the dynamics of cell turnover in the TM. We first confirmed prior descriptions of the proliferation and migration of cells over the pars tensa with a single EdU injection and chase (**Figure S6**) (Kakoi & Anniko 1997). Cycling cells were seen (1) near the handle of the malleus, (2) near the annulus, and (3) in the superior pars tensa at early time-points, and labeled cells had migrated to intermediate regions of the pars tensa by later time-points. We next wanted to identify which populations of cells were being replenished, and how rapidly they are turning over. We supplied mice with continuous EdU via their drinking water for three weeks and observed the accumulation of label, and then removed the source of EdU and observed the loss of label over four weeks (**Figure 3A**). EdU labeling over the malleus is dense by one week, and the area of EdU-positive cells expands outward until all of the keratinocytes have incorporated the label by three weeks (**Figure 3B**). The majority of cells incorporating EdU are keratinocytes, marked by expression of Krt5 (**Figure 3C**). Occasional fibroblasts and mucosal cells were also identified that had incorporated the EdU label during the three-week pulse. Thus, essentially all of the keratinocytes in the epidermis turn over in approximately three weeks, and the majority of other cellular populations in the TM are largely quiescent. The rapid turnover of the TM epidermis was again observed during the four-week chase period, when most of the EdU label in the keratinocytes dissipates (**Figure 3D-E**).

**Figure 3:**
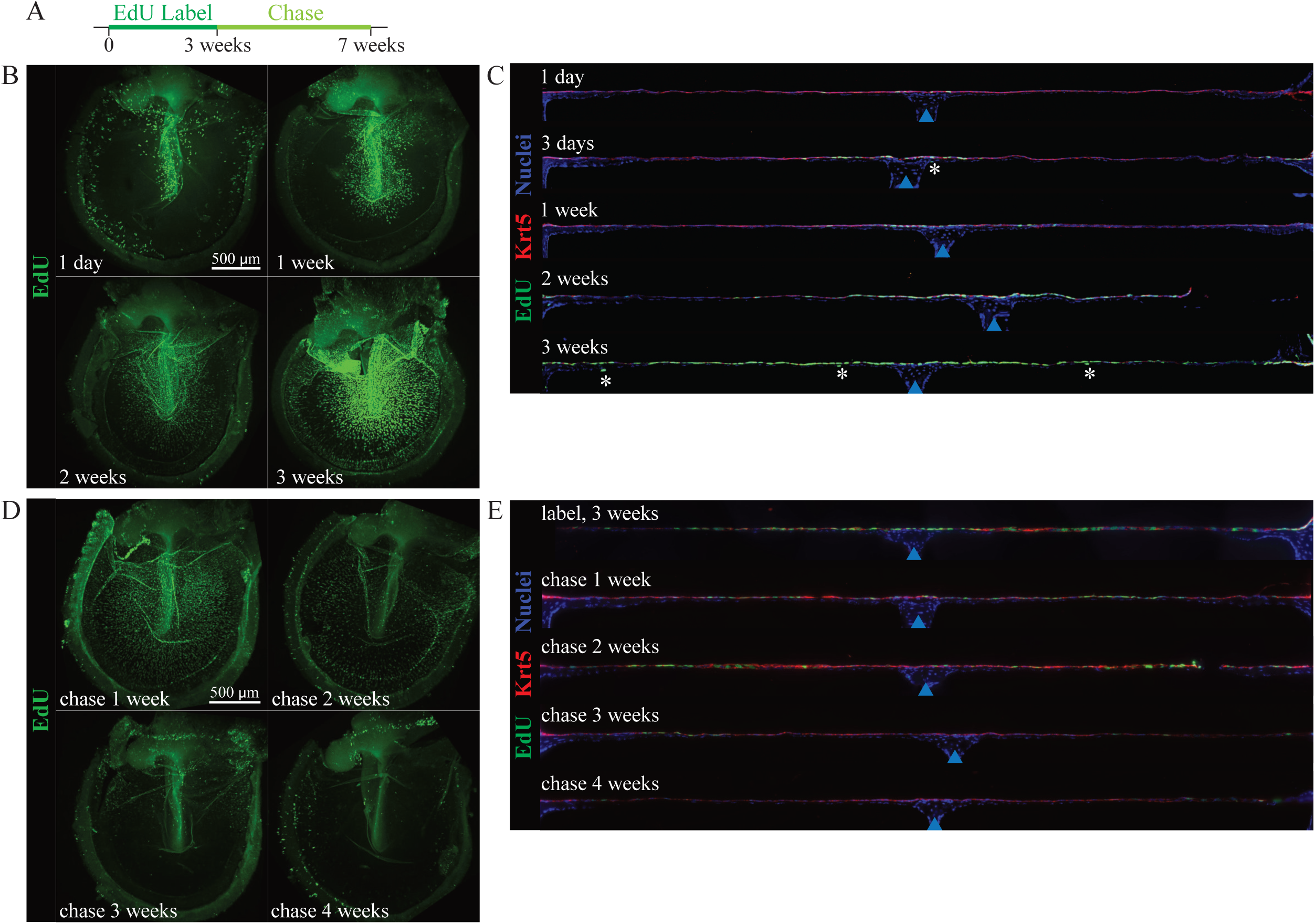
The murine TM epidermis turns over rapidly. (A) Timeline describing the pulse-chase experiment. Mice were exposed to EdU continuously for three weeks (pulse), and then the EdU source was removed and the label allowed to dilute for four weeks (chase). (B, D) Whole-mount TMs harvested at 1 day, 1 week, 2 weeks, and 3 weeks during the labeling period (B) or 1, 2 3, and 4 weeks during the chase period (D). TMs are stained for EdU (green). The scale bars indicating 500 μm apply to all four images in the same panel. (C, E) Full-width mid-malleus sections from TMs harvested at 1 day, 3 days, 1 week, 2 weeks, and 3 weeks during the labeling period (C) or 3 weeks during the labeling period and 1, 2, 3, and 4 weeks during the chase period. Sections are stained for EdU (green) and Krt5 (red). TMs were artificially straightened in Fiji (ImageJ); in each section, the tissue shown is approximately 1700 μm across. Asterisks indicate EdU+ cells that are not co-stained for Krt5. Blue arrows indicate the malleus.

### Live-cell imaging reveals TM keratinocyte migration patterns

To more deeply understand the migratory behavior of TM keratinocytes, we established techniques for time-lapse imaging of TMs cultured as explants. We employed two CreERT2 lines—Krt5-CreERT2 and Ki67-CreERT2—paired with two fluorescent reporters: mT/mG, in which labeled cells express membrane-localized EGFP, and R26R-Confetti, in which recombination leads to expression of one of four fluorescent proteins: RFP, YFP, mCFP, or nGFP (Rock et al. 2009; Basak et al. 2018; Muzumdar et al. 2007; Snippert et al. 2010). Krt5 is expressed diffusely throughout the TM (**Figure 1G**), so we anticipated extensive labeling with this line, whereas recombination induced in the Ki67-CreERT2 strains should be confined to the restricted proliferative zones.

In order to directly observe migratory behaviors of individual cells, a Krt5-CreERT2;mT/mG mouse was given a single dose of tamoxifen in order to label a subset of basal keratinocytes. The TM was harvested two days later, cultured as an explant, and imaged hourly over the next 24 hours. The cells were tracked with Imaris software. While radial movement was seen peripherally, the predominant direction of migration over the malleus was superior-to-inferior (**Figure 4A** and **Movie 1**). To observe the behavior of recently proliferated cells, we also performed live cell imaging on explants of Ki67-CreERT2;mT/mG mice. This revealed GFP predominantly arising from the pars flaccida and region near the malleus, and cell movements similar to those seen with the Krt5-CreERT2;mT/mG TM (**Figure 4B** and **Movie 2**). Cells of the pars tensa migrated with an average speed of 4.13 nm/second, while cells of the pars flaccida migrated with an average speed of 1.63 nm/second (**Figure 4C**). As a result, the average track displacement was 103.78 μm and 39.21 μm for cells of the pars tensa and pars flaccida, respectively (data not shown). Breaking the displacement into directional components, we again see that the pars tensa cells predominantly move superior-to-inferior (i.e., downward in the y-direction; **Figure 4D**). Thus the basal keratinocytes of the pars tensa exhibit directional lateral migration, and the basal keratinocytes of the pars flaccida do not. These populations may correspond to undifferentiated clusters 2 (*Fn1*+) and 1 (*Serpinb10*+), respectively, as cluster 2 has a transcriptional signature enriched for genes associated with a migratory phenotype (**Figures 4E** and **1M**).

**Figure 4:**
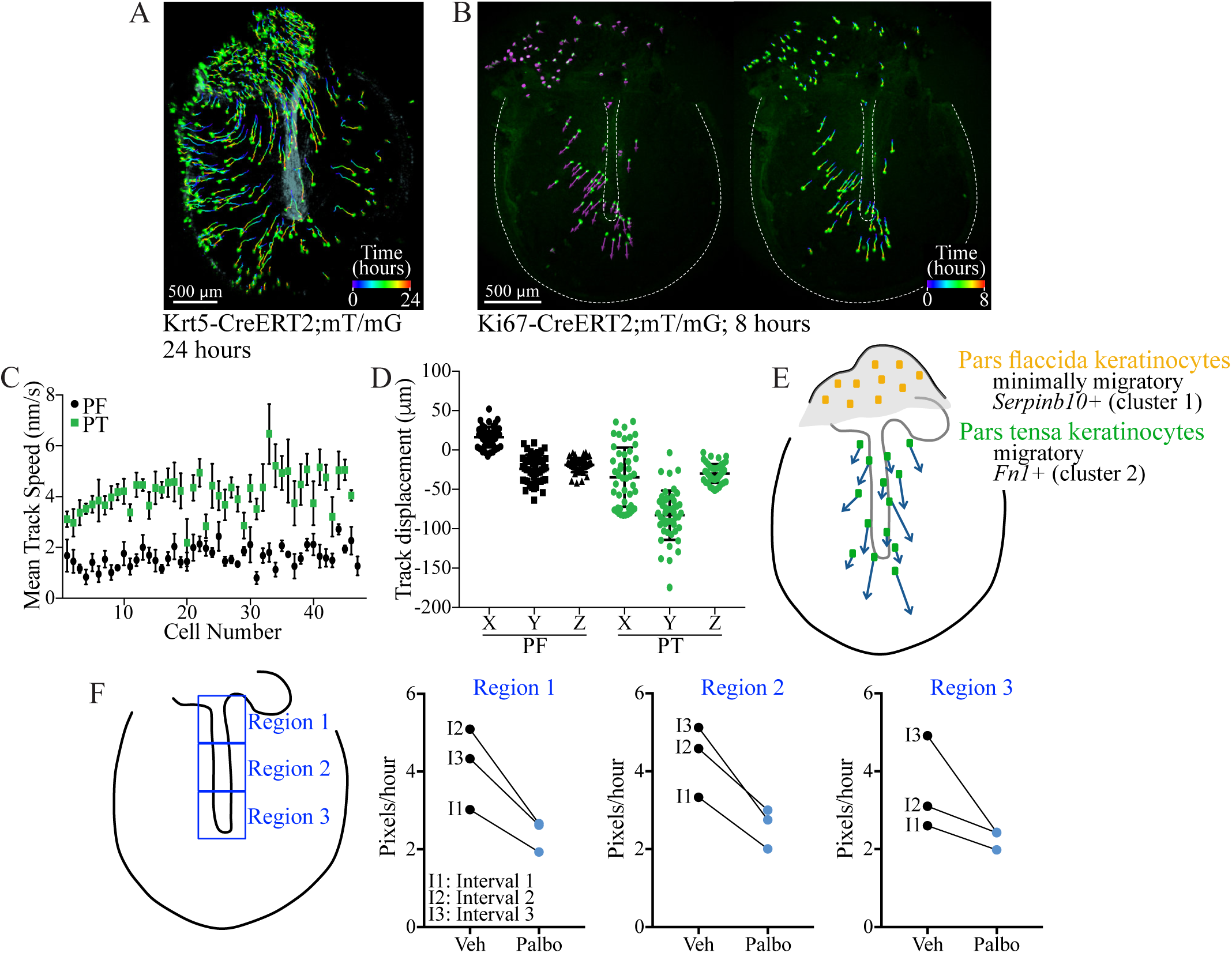
Live cell imaging of TM explants reveals prominent superior-to-inferior keratinocyte migration patterns. (A) TM from a Krt5-CreERT2;mT/mG mouse serially imaged hourly for 24 hours. Cells were identified and tracked using Imaris. Cells are shown at their final position, with their full track traced from time = 0 hours (purple) to 24 hours (red). (B) Whole-mount TM from a Ki67-CreERT2;mT/mG mouse imaged hourly for eight hours. On the left, cells are at their initial positions with vectors showing the directions in which they will move. On the right, cells are at their final positions with tracks showing their paths. (C) Mean speed of cells in the pars tensa (green squares) and pars flaccida (black circles). (D) X, Y, and Z components of the displacement. (E) Cartoon summarizing findings from the live cell imaging and scRNA-Seq experiments. (F) Speed of cell migration in TMs cultured in media with vehicle or 5 μM palbociclib. Results are shown for three distinct regions, as depicted in the cartoon on the left.

We next wanted to determine if the rate of keratinocyte migration is influenced by the generation of new cells in the tissue. To block cell proliferation, TMs were cultured in media with and without the Cdk inhibitor palbociclib and serially imaged four times over the course of 24 hours. In this experiment, imaged were aligned and cell tracks drawn manually. We calculated the speed of movement of each cell in the area over the malleus and found the cells in the palbociclib-treated TMs moved more slowly in all of the regions analyzed (**Figure 4F**). We stained TMs with cleaved caspase-3 to confirm that there was no increase in apoptosis with Cdk inhibitor treatment (data not shown). Thus, proliferation and migration in the TM appear to be partially coupled.

### Lineage tracing elucidates clonal architecture of TM keratinocytes

To further understand how proliferation and migration of TM keratinocytes cumulatively maintains the structure and function of the tissue, we next sought to determine the shape and location of clonal units of the TM epidermis. To visualize individual clones, we employed the Ki67-CreERT2;R26R-Confetti line. Mice were given a single dose of 250 mg/kg tamoxifen in order to label rare individual cells, and their TMs were evaluated two days to two months later (**Figures 5**).

**Figure 5:**
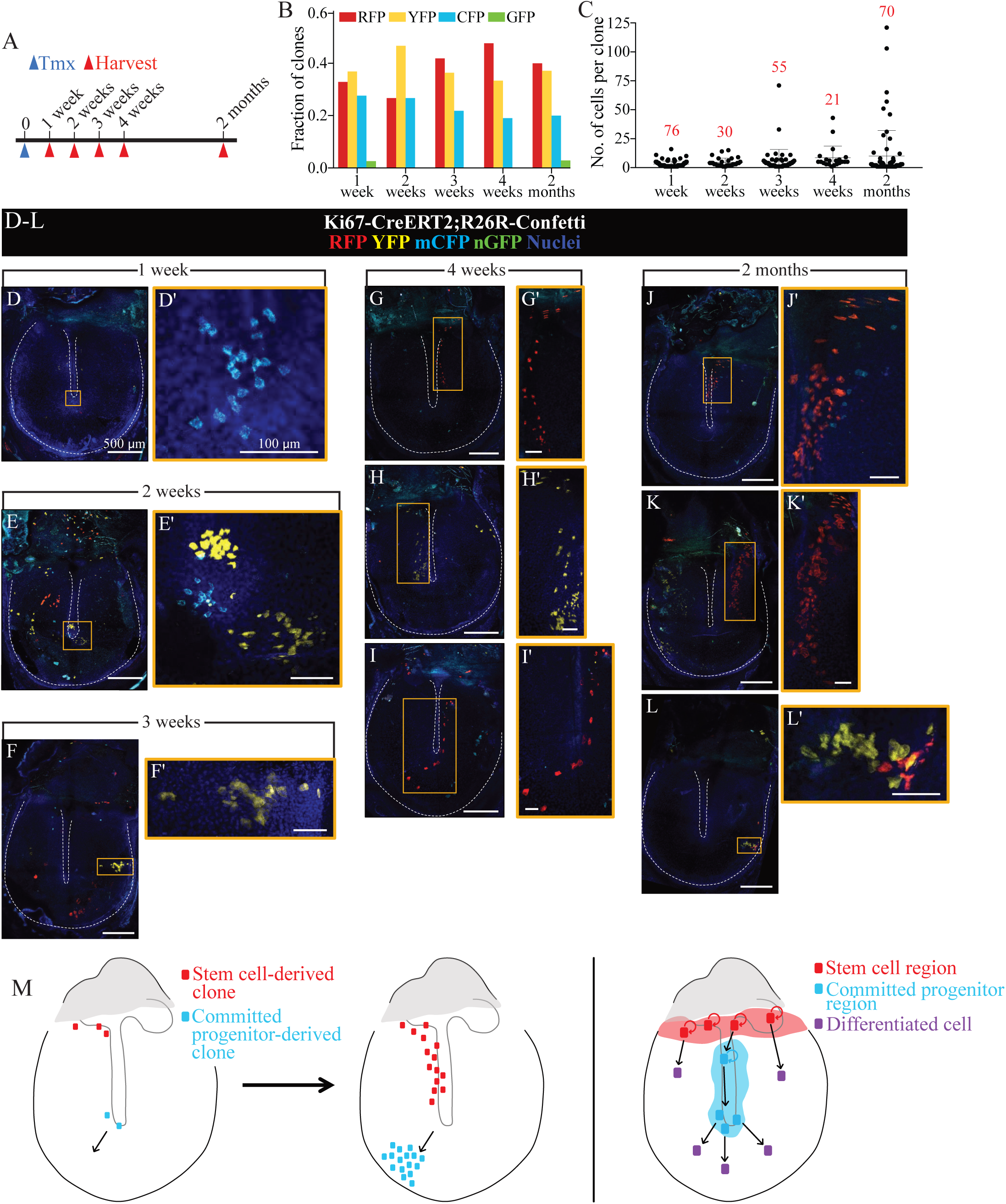
Single TM keratinocyte clones appear to arise from distinct stem cell and committed progenitor populations. (A) Experimental time-line. Mice were injected with 250 mg/kg tamoxifen at day 0 and TMs were harvested weeks to months later. (B) Fraction of clones expressing the different fluorescent proteins at each time-point. (C) Number of cells per clone identified at each indicated time-point. The total number of clones quantified at each time-point is indicated above the data. (D-L) Examples of clones captured at the indicated time-points. In each pair of images, the entire whole-mount TM is shown on the left with the annulus and malleus outlined by a white dotted line, the magnified area shown with a yellow box, and the scale bar indicating 500 μm; on the right, designated with a prime, are more magnified images of clones, and all of the scale bars in these images indicate 100 μm. (M) Cartoon depicting our model for the location of stem cells (SCs) and committed progenitors (CPs) of the murine TM epidermis based on data from the Ki67-CreERT2;R26R-Confetti lineage trace. We posit that SCs reside in the superior portion of the pars tensa, near the border with the pars flaccida, and that CPs proliferate over the malleus.

Two days after tamoxifen injection, 0-10 fluorescent cells per pars tensa were seen (data not shown). In the 39 TMs evaluated between one week and two months following recombination, 252 clones were confidently identified. Of these, only four expressed nGFP; 94 were RFP+, 95 were YFP+, and 59 were mCFP+, with similar distributions at each individual time-point (**Figure 5B**). The number of cells in each clone was counted, and the majority of clones at every time-point contained ten or fewer cells (**Figure 5C**). However, the distribution of clone sizes expands at later time-points: between one and two weeks, the maximum clone size is 16 cells; between three and four weeks, it is 71 cells; and at two months, it is 121 cells. This scaling behavior has been described in many other tissues, including testis, interfollicular epidermis, and intestine, and is consistent with neutral drift dynamics in the SC pool (Klein et al. 2010; Mascré et al. 2012; Lopez-Garcia et al. 2010).

The shape and location of the larger clones was further informative. The cells were not contiguous in the clones over the pars tensa, indicating that cells from multiple progenitors are rapidly interweaved as they migrate across the TM (**Figure 5D-L**). At one to two weeks, we captured several relatively circular clones near the umbo of the malleus, which was previously identified as a major site of cell turnover (**Figure 5D-E**) (Knutsson et al. 2011; Liew et al. 2018). At later time-points, we never observed a streak radiating exclusively from the malleus to the annulus. These patterns are consistent with a model whereby proliferating cells around the malleus are CPs and subsequently migrate away from the umbo and towards the EAC during the 21-day turnover period of the TM (**Figure 5M**).

Another set of informative clones, observed in TMs four weeks or two months after recombination, were elongated along the handle of the malleus or in streaks extending supero-inferiorly along the pars tensa, but always connecting back to the border with the pars flaccida (**Figure 5G-K**). The morphology of these clones is consistent with the superior-to-inferior pattern of migration observed in the time-lapse studies. Furthermore, most of these clones include cells in the upper pars tensa, near the border with the pars flaccida, which is a known proliferative region. Due to the presence of clones emanating from this region at later time-points, we posit that the superior region of the TM near the junction of the pars flaccida and pars tensa is the location of the long-term repopulating SCs of the tissue (**Figure 5M**).

To test for possible rare contributions from long term repopulating SCs existing along the malleus, we assayed for the emergence of clonal architecture in densely labeled TMs. Maximal labeling was induced in Krt5-CreERT2;R26R-Confetti mice via five days of two mg/day tamoxifen administration. TMs were evaluated three days and one, three, and six months following recombination (**Figures 6 and S8**). At three days, keratinocytes throughout the TM are labeled, and cells expressing RFP, YFP, and mCFP are randomly assorted. Consistent with prior observations, very few nGFP-expressing cells are seen. At six months, the entire TM epidermis remains fluorescent, indicating that the long term repopulating SCs maintaining this tissue layer are indeed Krt5+. Over the pars flaccida, color blocks rapidly emerge, consistent with prior observations in normal interfollicular epidermis. This also is consistent with the relative lack of migratory behavior in the cells of the pars flaccida.

**Figure 6:**
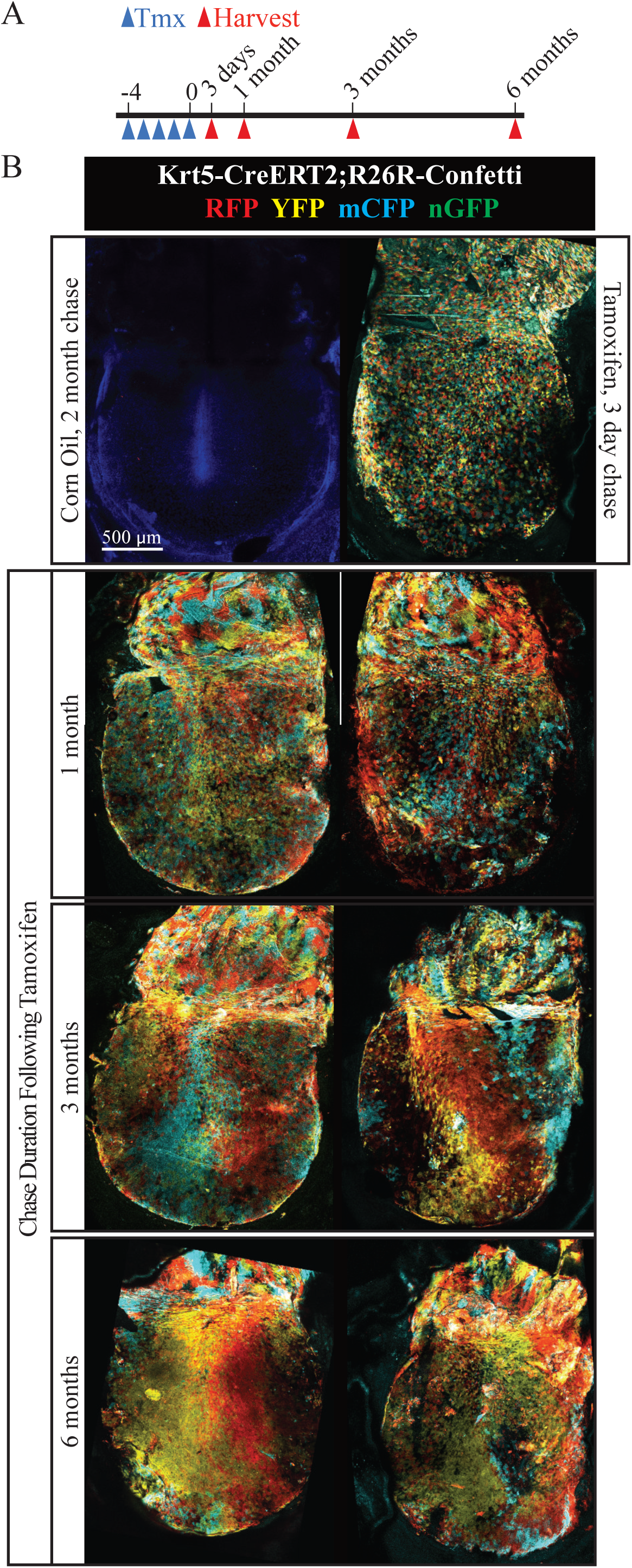
Clonal drift in densely labeled TMs. (A) Experimental time-line. Krt5-CreERT2;R26R-Confetti mice were given 2 mg/day tamoxifen for 5 days and then sacrificed 3 days to 6 months later. (B) Representative whole-mount TMs from the indicated time-points. A TM from a control animal dosed with corn oil for 5 days and sacrificed two months later is included at the top left. The remaining TMs are from mice given 5 days of tamoxifen. The scale bar indicating 500 μm applies to all images.

In contrast, color blocks did not emerge at early time-points in the pars tensa, consistent with the active lateral migratory behavior and cell mixing of pars tensa keratinocytes. However, at time-points after multiple cycles of complete turnover of the TM, streaks of color formed. Such coarsening is considered a hallmark of stochastic stem cell turnover (Klein & Simons 2011). Furthermore, in each case, the streaks were oriented supero-inferiorly across the TM and always connected back to the border with the pars flaccida, and not necessarily to the region over the malleus. To quantify the progression towards tissue-wide oligoclonality, a grid was superimposed over the TM and each 5,000-μm^2^ box was scored as containing cells of one to three colors (**Figure S8**). At 3 days after inducing maximal recombination, the average percentage of boxes with mCFP, YFP, and RFP expression was over 95%. At 1 month, this remained stable at 92%, consistent with the rapid mixing of cell progeny due to migration. At 3 and 6 months, the percentage of boxes with all three fluorescent proteins drops sharply, and regions with 2 or 1 color increase, consistent with the emergence of color blocks apparent by visual inspection.

Because we observe dense labeling even 6 months following induction of the lineage trace, we conclude that a subset of Krt5+ cells act as the long term SC of this tissue. Further, the architecture of the clones is consistent with a model whereby long term repopulating SCs arise exclusively from the region adjacent to the pars flaccida, and their CP and differentiated progeny migrate over the pars tensa in order to maintain a thin, vibratory surface free of cellular and exogenous debris.

### TM fibroblasts provide niche signals that promote keratinocyte turnover

The restricted pattern of proliferation in the TM suggested niche signals restricted to these proliferative zones. We next aimed to identify molecular signals that are required for proliferation of TM keratinocytes. When cultured as explants in non-supplemented media, TMs maintain regionally restricted proliferation predominantly near the malleus and in the pars flaccida. We utilized this system to screen 74 compounds with defined molecular targets for reduction in EdU incorporation (**Table S3**). The compound library was designed to include only molecular targets whose expression was confirmed in the murine TM scRNA-Seq dataset, and it was predominantly comprised of cell surface receptors. TMs were cultured in media containing vehicle (n = 26) or 40 μM of a compound (n = 2 per compound) for 48 hours then pulsed with EdU for two hours. They were then processed and imaged, and the number of EdU-positive cells in a 1.2 mm × 1.6 mm region of the pars tensa were counted (**Figure 7A-B**). Compounds targeting Cdks (SNS-032 and Palbociclib) or Aurka (Tozasertib) were included as positive controls, and they ablated proliferation as expected. Targets for which every compound tested ablated proliferation included: Bmpr (K02288, LDN-193189), Fak (PF-573228, TAE226), Fgfr2/3 (AZD4547), Ikk2 (Tpca-1, IMD 0354), Pdgfr (CP-673451, Crenolanib), Ppar (GW9662, FH535), S1pr/Sphk1 (Ozanimod, PF-543), and Trpv4 (GSK2193874).

**Figure 7:**
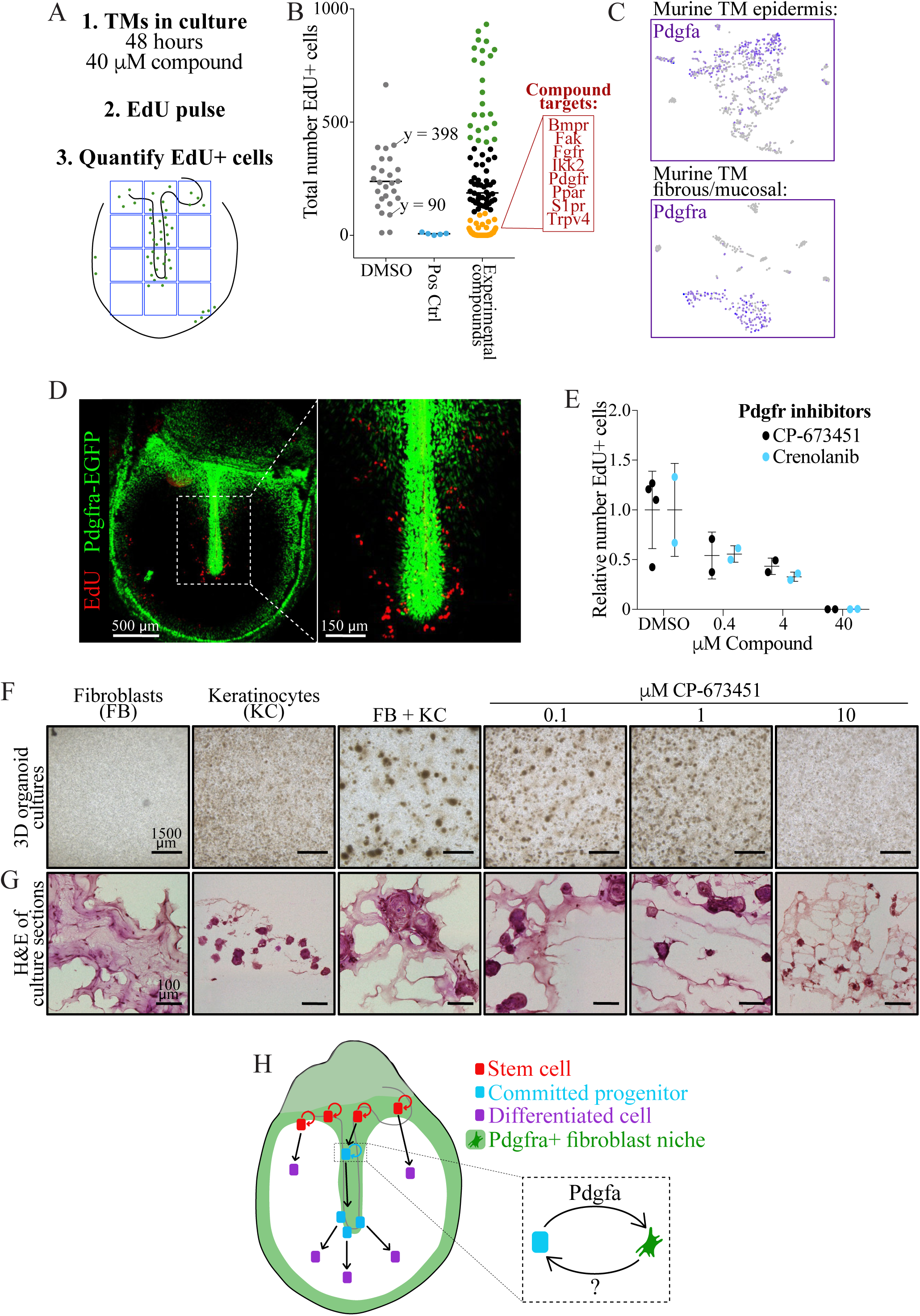
Pdgfra signaling in fibroblasts supports turnover of mouse and human TM keratinocytes. (A) Approach for screening compounds in TM explant cultures. (B) Summary of the screen results. The numbers of EdU+ cells in consistent 1,200 μm × 1,600 μm areas of the TMs are plotted. Results from DMSO-treated TMs are shown in grey and positive control TMs (treated with inhibitors of Cdk (SNS-032, Palbociclib) or Aurka (Tozasertib)) in blue. Of the TMs treated with one of 71 experimental compounds, data points are colored in black if the EdU+ cell count was in the main DMSO-treated range (90 – 398), in green if the count was greater than 398, and in gold if the count was less than 90. In the red box are targets for which every compound tested fell in the gold range. (C) tSNE plots showing expression of Pdgfa ligand in the murine TM epidermis (top) and of Pdgfra receptor in the murine TM fibrous/mucosal fraction (bottom). (D) Whole-mount TM from a Pdgfra-EGFP mouse injected with EdU and sacrificed 24 hours later. EdU is stained in red. (E) Relative number of EdU+ cells in TM explants cultured for 48 hours in vehicle or the indicated concentrations of Pdgfra inhibitor CP-673451 or Crenolanib. (F) Images of three-dimensional organoid cultures of fibroblasts and keratinocytes from human TMs, seeded separately or together in co-culture. Fibroblasts and keratinocytes co-cultured were also treated with the indicated concentrations of CP-673451. (G) H&E stained sections of the organoid cultures. (H) Cartoon summarizing our findings. Pdgfra+ fibroblasts create a niche (green) supporting proliferation of keratinocyte stem cells (red) and committed progenitors (blue), while keratinocytes not in proximity to this niche differentiate (purple).

We queried the scRNA-Seq datasets to determine where these targets were expressed and noted that, while Pdgfra is confined to the mesenchyme, the ligand Pdgfa is expressed by the undifferentiated keratinocyte clusters, including cycling cells (**Figure 7C**). Immunofluorescence for Vim demonstrated that the mesenchyme is confined to known regions of TM proliferation (**Figure S3**). To more specifically identify the anatomic location of the Pdgfra positive cells and their proximity to cycling keratinocytes, we administered a single dose of EdU to a Pdgfra-EGFP reporter mouse, and noted a co-localization of Pdgfra-expressing cells and proliferating cells in similar areas of the TM (**Figure 7D**) (Hamilton et al. 2003). Because the screen was done using high compound concentrations, we treated explants with multiple concentrations of two Pdgfr inhibitors and saw a concentration-dependent decrease in cell proliferation (**Figure 7E**). Thus, Pdgfr inhibition blocks proliferation of TM keratinocytes.

Lastly, we asked if primary Pdgfra+ fibroblasts from a human TM support the growth of primary human TM keratinocytes. These two cell populations were isolated from surgical TM samples, and either 8,000 cells of a single type or a 50:50 mix of keratinocytes and fibroblasts were plated in an organoid assay (**Figure 7F-H**) (Frank et al. 2016). When plated in solitary cultures, the fibroblasts do not form any clear multicellular structures and the keratinocytes form keratin pearls—spherical clusters of concentric keratinocytes often identified in squamous cell carcinomas. In co-culture, however, larger colonies comprised of a keratinocyte core and fibroblast periphery are formed. Treatment of the co-cultures with the Pdgfr inhibitor CP-673451 reverted the phenotype of these colonies (**Figure 7F-H**). Therefore, we conclude that the keratinocyte-stromal interaction mediated by Pdgfr signaling is likely conserved in the human TM.

## Discussion

### The TM is a unique epidermal site

Our study elucidates the proliferative, migratory, and clonal dynamics of keratinocytes of the tympanic membrane (TM). The TM conducts sound waves from the external auditory canal (EAC) to the ossicular chain, which in turn transmits them to the inner ear where they are converted to neural signals. It is comprised of an outer epidermal layer continuous with that of the EAC, an inner mucosal layer continuous with that of the middle ear, and a middle fibrous layer. The epidermis is in many ways similar to epidermis at other anatomical sites; for example, it is a barrier against infectious agents. But, its position at the blind end of the ear canal necessitates an additional physiological function: clearing the canal of squamous debris and foreign objects. To achieve this, key zones of proliferation are in the center and superior-most regions of the pars tensa, as confirmed here by EdU labeling, and keratinocytes migrate inferiorly and radially outward, as we have directly visualized with live cell imaging. This arrangement is similar to what is seen in re-epithelialization following wounding, where a proliferative region exists behind a migrating front (Park et al. 2017). Until now, to our knowledge this has not been recognized as a phenomenon that occurs under normal physiological circumstances.

Tissue thickness is a key functional property of the epidermis that differs by anatomic location. Across the TM alone, the epidermis varies in thickness from a single cell to roughly five cells. The epidermis of the pars flaccida is thicker and thus more stratified than the epidermis of the pars tensa. In our single-cell RNA sequencing (scRNA-Seq) data, we captured cells spanning the differentiation spectrum. Staining for *Flg2*, a marker usually expressed by differentiated keratinocytes of the stratum granulosum, showed markedly more differentiation in the pars flaccida, as would be expected. In much of the pars tensa, there is no cell stratification, and concomitantly few terminally differentiated keratinocytes were seen. However, there were occasional *Flg2*+ keratinocytes in the plane of the epidermal monolayer of the pars tensa. This is in contrast to what occurs in skin elsewhere, where differentiation and stratification are obligately coupled (Lechler & Fuchs 2005; Clayton et al. 2007). How this differentiation is induced independent of exit from the basal layer, and whether it is linked to the lateral migration of keratinocytes instead of to stratification, are questions for further study.

### Clonal dynamics in the TM

The atypical arrangement and migratory behavior of the epidermis of the TM reveals new clonal dynamics of epidermal keratinocytes. The lineage tracing analyses show the net results of keratinocyte proliferation and migration. In regards to maintenance of the TM epidermal stem cell (SC) pool, both Ki67-CreERT2 and Krt5-CreERT2 traces are qualitatively consistent with a model of population asymmetry and neutral drift, in which some SCs are lost over time and others divide symmetrically to maintain the SC pool (Klein & Simons 2011). Quantification of single clones from Ki67-CreERT2;R26R-Confetti mice demonstrated widening of the distribution of clone sizes, also known as scaling behavior, over time. In traced Krt5-CreERT2;R26R-Confetti samples, the emergence of more distinctive and larger color blocks over time also supports this model.

A key challenge in interpreting these data is that the SC niche of the TM is not well defined. We know generally where the proliferative zones are, but we also know that most of the TM keratinocytes become proliferative following injury (Chari et al. 2019). Together, the live imaging and lineage tracing studies support the hypothesis that the cells that maintain the TM epidermis long-term reside in the superior region of the pars tensa or potentially in the pars flaccida. In the live imaging analysis, the direction of migration is largely superior-to-inferior. At two months in the Ki67-CreERT2 studies, a relatively late time-point, none of the larger clones documented included cells near the umbo, but several included cells in the superior region of the TM. Finally, at three months in the Krt5-CreERT2 experiments, the predominant orientation of distinctive color blocks is vertical. All of these findings are consistent with a more stable progenitor niche in the superior pars tensa. The proliferating cells over the malleus behave more like committed progenitors (CPs), making a limited contribution to the tissue before migrating into the ear canal. Future work to explore these hypotheses will include selective labeling of different regions of the TM, as well as *in vivo* time-lapse analysis.

Previous data from other areas of mouse epidermis has yielded conflicting results depending on the particular methodology used to distinguish between neutrally competing stem cells and a hierarchical model (Fuchs 2016). The spatial separation of keratinocyte SCs and CPs within the TM presents the clearest evidence we know yet of the capacity of keratinocytes to exist in a clonal hierarchy. It should be noted that the fact that this arrangement exists within the TM does not necessarily argue directly for such an arrangement in other areas of the epidermis. However, it does argue that given the proper signals such an arrangement may arise.

### Lateral keratinocyte migration establishes the clonal hierarchy

Although it appears that in uninjured TM the keratinocytes of the majority of the pars tensa do not have long-term repopulation activity, they may act as facultative SCs in response to injury. Indeed, we and others have previously demonstrated a massive proliferative response in keratinocytes throughout the pars tensa following perforation (Santa Maria et al. 2010; Chari et al. 2019). One possibility is that the majority of the keratinocytes of the pars tensa are capable of long-term repopulation activity, but lack the niche signals to execute that program. This model may help explain the pattern of proliferation observed in the different parts of the TM. If the proper niche signals to maintain proliferation come predominantly from stromal cells (perhaps Pdgfra+ cells), then, as the progeny of the long term repopulating cells migrate inferiorly, if they happen to be in proximity to those niche signals—such as over the malleus—they will continue to proliferate. If they are not in proximity to those signals—as in over the body of the pars tensa—then they may continue to migrate but would cease proliferation. Because the proliferative cells over the malleus would inexorably move away from the malleus, they would be removed from the proximity to the niche signals and therefore stop proliferating. In this way, the migration itself may confer the hierarchical behavior, rather than an enforced cell-autonomous differentiation program.

One outstanding question is how the direction of migration of the TM keratinocytes is enforced. Although we do not yet have direct evidence to validate a mechanism, we suspect that it may be related to inherent properties of the pars tensa. Previous work has shown that asymmetries in the tension of structures on which keratinocytes are grown may confer directional migration (Lü et al. 2013; Lü et al. 2016). Under this model, when progeny of TM SCs transition inferiorly onto the body of the pars tensa, they experience vectors of force that were not present in the uppermost pars tensa or pars flaccida and begin to migrate along those vectors. An additional and not mutually exclusive possibility is that arrangement of collagen fibers in the pars tensa permits directional migration in a way that the extracellular matrix of the pars flaccida does not. Deep to the basement membrane there is a highly organized network of radial collagen fibers that may approximate some of the migratory patterns of the keratinocytes (Fay et al. 2006). These hypotheses bear further experimental validation.

### Niche signals that maintain keratinocyte proliferation

The stromal signals required for the maintenance of keratinocyte proliferation within uninjured epidermis are not well defined. In order to maintain primary keratinocytes in culture, media formulations require a rich mix of growth factors that may replicate aspects of wound healing rather than physiologic maintenance. Further, organotypic models of stratified squamous epithelia almost always require fibroblasts as “feeders” to maintain the keratinocytes in a healthy, renewing state (Oh et al. 2013). Despite the well-established necessity of stroma in supporting keratinocytes, the exact mechanisms by which this activity is executed remain cryptic. We propose Pdgf signaling as one critical component of this crosstalk.

Because basal keratinocytes of the TM express Pdgf and mesenchymal cells express Pdgfr, we propose that keratinocytes are sending trophic signals to the mesenchyme. Given that inhibition of Pdgfr ablates proliferation of keratinocytes, this argues for signals then being sent back from the mesenchyme to the keratinocytes. We do not yet know the identity of these signals, but they may be diffusible factors, membrane bound factors, extracellular matrix components, or possibly even mechanical forces. Further studies will help clarify these mechanisms. Because of the limited zone of proliferation within the pars tensa, the TM may be an ideal model system to help understand the mechanisms by which keratinocyte-stromal interactions maintain keratinocyte proliferation. With a better understanding of these interactions, we may then extend these findings to epidermis of other parts of the body, as well as other squamous epithelia.

### Implications for human disease

Lastly, TM physiology is further important to understand because it is deranged in pathological conditions specific to the TM. Abnormal migration has been documented in keratosis obturans, a condition in which desquamated keratin builds up in the ear canal, and myringitis granulosa, a chronic inflammatory condition of the lateral TM (Corbridge et al. 1996; Soucek & Michaels 1993; Makino et al. 1988). TM cell proliferation and migration have also been implicated in closure of TM perforations, which occurs spontaneously in ∼90% of instances (Lou et al. 2012; Lou 2012; Chari et al. 2019). In the remaining cases, chronic perforations develop and can result in hearing impairment, infections, and drainage. In all of these pathologies, advances in care have been hampered by lack of understanding of the normal physiology of the tissue.

Another example is cholesteatoma, an abnormal growth of TM keratinocytes that invades the middle ear space and can destroy structures in and surrounding the ear (Kuo et al. 2015). The underlying pathogenesis of this condition is not well understood, but this study suggests a potential mechanism. The vast majority of spontaneous acquired cholesteatomas occur as a consequence of retraction of the pars flaccida or superior pars tensa. If there is minimal lateral migratory activity of the pars flaccida, then, following retraction, progeny of proliferating cells may simply stratify and accumulate within the retraction pocket. In contrast, because there is minimal stratification and rapid lateral migration of the keratinocytes of the pars tensa, retraction in that area would not give rise to the accumulation of squamous debris. The findings herein thus provide a basis for deepening understanding of cholesteatoma and other conditions of the TM, as well as basic epidermal biology.

## Author Contributions

Conceptualization, A.D.T. and S.M.F.; Methodology, A.D.T., S.M.F., and J.C.; Formal Analysis, S.M.F., K.S.Y., J.C., and K.P.L.; Investigation, S.M.F., J.C., L.E.B.; Resources, A.D.T., J.A.B., and J.B.S.; Writing, A.D.T. and S.M.F.; Visualization, A.D.T. and S.M.F.; Supervision, A.D.T. and J.B.S.

## Acknowledgements

We thank Ophir D. Klein, Diana J. Laird, and Tien Peng for helpful discussions and reading of the manuscript (O.D.K. and D.J.L.); Bruce Wang and Licia Selleri for use of their microscopes; the Biological Imaging Development Center, Broad Center Microscopy Core, and Institute for Human Genetics for their expertise and equipment. Also, we thank our funding sources: American Otological Society (S.M.F. and A.D.T.) and the Hearing Research Inc. (A.D.T.).

## STAR Methods

Further information and requests for resources and reagents should be directed to and will be fulfilled by the Lead Contact, Aaron D. Tward.

## EXPERIMENTAL MODEL AND SUBJECT DETAILS

### Animals

Mice were maintained and experiments conducted following the guidelines of the Institutional Animal Care and Use Committee at the University of California, San Francisco (approval number AN165495). All experiments using wild type mice were done with strain FVB/NJ. All Cre driver lines (Ki67-CreERT2, Ki67CreERT2, and Krt19-CreERT) and floxed reporter lines (R26R-Confetti and mT/mG) were received and maintained on a C57BL/6 background. Pdgfra-EGFP mice were received and maintained on a FVB/NJ background. In most cases, adult animals—both male and female—between 6 and 12 weeks of age were used. Additional details are included in the Methods Details section below.

### Primary human tissue samples for scRNA-Seq and organoid experiments

Normal TM tissue was obtained from surgical procedures at University of California, San Francisco under approval of the IRB (approval number 16-19811) and with informed consent from the patients. For scRNA-Seq, TM tissue was removed from a 34-year-old female as part of a surgical approach for unrelated tumor adjacent to the brain. Fibroblasts and keratinocytes used in the organoid assay were derived from undiseased TM tissue from patients undergoing surgery for cholesteatoma in the pars flaccida region of the TM. Fibroblasts were derived from tissue from a 55-year-old male, and keratinocytes were derived from tissue from a 27-year-old female.

## METHODS DETAILS

### TM dissociations

#### Murine

TMs were dissected from 19 6-week-old FVB/NJ mice (10 female, 9 male). They were incubated in dispase (Corning) at 37°C for five minutes and then mechanically separated into epidermal and fibrous/mucosal fractions. The epidermal tissue was dissociated in TrypLE (Life Technologies) and the fibrous/mucosal tissue was dissociated in 0.2 mg/mL collagenase P (Sigma-Aldrich) and 5 μg/mL DNAse. Both dissociations were done at 37°C for 10 minutes, with trituration every five minutes. Cells were passed through 40 μm strainers, collected by centrifugation, and subjected to removal of dead cells (Miltenyi Biotec). They were resuspended at 1,000 cells/μL in 0.04% BSA in phosphate buffered saline (PBS), and 25,000 cells were loaded for single cell capture.

#### Human

Prior to dissociation, the TM was mechanically separated into an outer epidermal portion and an inner fibrous/mucosal portion. A scalpel was used to cut the tissue into pieces less than two mm. The outer TM was dissociated in 1 mg/mL collagenase P and 5 μg/mL DNAse at 37°C for one hour, with trituration every 20 minutes. The inner TM was incubated in dispase at 37°C for 30 minutes and then transferred to collagenase P and DNAse for an additional hour, with trituration every 20 minutes. Cells were passed through 40 μm strainers and collected by centrifugation. Red blood cells were lysed (Alfa Aesar J62990), and then dead cells were removed (Miltenyi Biotec). Cells were resuspended at 1,000 cells/μL in 0.04% BSA in PBS, and 25,000 cells were loaded for single cell capture.

### scRNA-Seq analysis

Isolated cells were run on the Chromium Controller (10X Genomics) with Single Cell 3’ Reagent Kit v2 and the generated libraries were sequenced on Illumina HiSeq 4000. Human data was aligned to hg19 and mouse to mm10. Data was run through CellRanger 2.0.0 (10x Genomics) and then analyzed via R primarily through single cell analysis package Seurat (R Core Team 2018; Satija et al. 2015). Unwanted sources of variation were processed through removal of cells with more than 7,000 genes and regression was done via UMIs. All genes expressed in fewer than three cells were removed, as were all cells that expressed fewer than 200 genes. The matrices of data were log normalized in a sparse data matrix and principal component analysis was performed and the first 10 components were used to cluster the cells by Louvain clustering implemented in Seurat. tSNE plots were independently generated to aid in 2D representation of multidimensional data.

### Immunofluorescence (IF)

Whole-mount TMs were either dissected en bloc, or the auditory bulla was isolated and the TM dissected out following fixation and decalcification. Following dissection, TMs were fixed in 4% paraformaldehyde (PFA) at 4°C for two hours (en bloc) or overnight (auditory bullae). They were decalcified in 5% EDTA overnight (en bloc) or for one to three days with daily solution changes (auditory bullae). TMs were permeabilized in PBS with 1% Triton X-100 for two hours at room temperature (RT), blocked in PBS with 1% Triton X-100 and 10% fetal bovine serum (FBS; blocking buffer) for two hours at RT, and incubated in primary antibody diluted in blocking buffer at 4°C overnight. They were washed and then incubated in secondary antibody in blocking buffer for one hour at RT. They were again washed and, if nuclear stain was desired, incubated in Hoechst dye diluted 1:2,000 in PBS for 30 minutes. They underwent final washed and were then mounted. Whole-mount TMs were imaged on a Nikon AZ100 macro confocal microscope. Images were processed using Fiji (ImageJ) software (Schindelin et al. 2012). Whole-mounts are shown as maximum projections of z-stacks.

To prepare paraffin sections, auditory bullae were fixed and decalcified as above, dehydrated and embedded in paraffin, and then sectioned at 7-μm thickness. Staining was done as previously described(Hu et al. 2017). Briefly, paraffin was dissolved in Histo-Clear II (Electron Microscopy Sciences) and then the tissue was rehydrated. Slides were sub-boiled in citrate buffer (10 mM citric acid, 2 mM EDTA, 0.05% Tween-20, pH 6.2) for ten minutes in a microwave to retrieve antigens. Samples were permeabilized and blocked at room temperature for one hour in Animal-Free Blocker (Vector Labs SP-5030) supplemented with 1% Triton X-100 and 2% normal goat serum (Cell Signaling Technology). They were incubated in primary antibody diluted in blocking buffer at 4°C overnight. The following day, slides are washed and then incubated in secondary antibody at RT for one hour. They were again washed and mounted with ProLong Gold Antifade Mountant with DAPI (Life Technologies). Images were acquired on a Leica DM6 B microscope and processed using Fiji (ImageJ) software.

### *In situ* hybridization (ISH)

Prior to dissection of TMs to be used for *in situ* hybridization (ISH), mice were euthanized with CO_2_ and thoracotomy, and then their vasculature was perfused with RNase-free PBS followed by 10% normal buffered formalin (NBF; sections) or 4% PFA (whole-mounts) to fix the tissue *in situ*. For sections, auditory bullae were then isolated, incubated in NBF at RT for 24 hours, then decalcified, paraffin embedded, and sectioned as described for IF. The RNAscope Multiplex Fluorescent Reagent Kit v2 (Advanced Cell Diagnostics) protocol was followed, with the following specifications: manual target retrieval was done for 15 minutes and digestion with Protease Plus for 30 minutes. Images were acquired on a Leica DM6 B microscope and processed using Fiji (ImageJ) software.

For whole-mount TMs, mice younger than 5-weeks-old were used so that the TM could be dissected out in a bony ring without the need for decalcification. A protocol for staining of whole-mount zebrafish embryos was followed (Gross-Thebing et al. 2014). Briefly, the TMs were fixed in 4% PFA diluted in RNase-free PBS for 6 hours at RT, then washed in PBT (RNase-free PBS with 0.1% Tween 20), dehydrated in increasing concentrations of methanol, then stored in 100% methanol at −20°C overnight or for several days. The methanol was removed and the TMs air-dried, then incubated in Protease III at RT for 20 minutes, washed, and probes hybridized at 50°C overnight. The following day, the TMs were washed in 0.2X SSCT, fixed in 4% PFA at RT for 10 minutes, and again washed. The signal was then amplified and developed following the steps in the RNAscope Multiplex Fluorescent Reagent Kit v2 protocol. Images were acquired on a Nikon AZ100 macro confocal microscope and processed with Fiji (ImageJ).

### EdU administration and detection

For injection, EdU (Carbosynth Limited) was resuspended at 5 mg/mL in saline and passed through a 0.2 μm filter. One mg was given by intraperitoneal (IP) injection. For continuous administration, mice were given an IP injection at the start of the experiment and then provided with EdU in the water, resuspended at 0.5 mg/mL with 1% sucrose and passed through a 0.2 μm filter. Water bottles were shielded from light and the water supply changed every three days. Samples were dissected and processed as for IF. For EdU detection, the Click-iT EdU Alexa Fluor 488 Imaging Kit (ThermoFisher Scientific) was used. For combined EdU and protein detection, the IF protocol was followed after EdU detection, beginning at the blocking step.

### Live-cell imaging

To label TMs for live cell imaging, recombination was induced in Ki67-CreERT2;mT/mG or Krt5-CreERT2;mT/mG mice with a single IP injection of 250 mg/kg or 1 mg tamoxifen, respectively. TMs were harvested en bloc and cultured floating on Advanced DMEM (Life Technologies 12491023) supplemented with 1% penicillin/streptomycin (Fisher Scientific). When relevant, compounds or vehicle were added to the media. Live-cell imaging was done on a Nikon AZ100 macro confocal microscope in an incubated chamber maintained at 37°C with 5% CO_2_ and 95% humidity. Z-stacks were acquired approximately every hour. Cells were identified and tracked using Imaris software.

### Lineage tracing with the R26R-Confetti reporter

For minimal labeling, Ki67-CreERT2;R26R-Confetti mice were given a single IP injection of 250 mg/kg tamoxifen. For maximal labeling, Krt5-CreERT2;R26R-Confetti mice were given 2 mg tamoxifen per day for 5 consecutive days. At the specified time-points, auditory bullae were harvested, fixed in 4% PFA at 4°C for 4 hours, decalcified in 5% EDTA at 4°C overnight, and then the bone was resected to leave the TM in a bony ring. Krt5-CreERT2;R26R-Confetti TMs were directly mounted for imaging. Prior to mounting, Ki67-CreERT2;R26R-Confetti TMs were permeabilized in 0.5% Triton X-100 in PBS at RT for 2 hours, and nuclei were stained with Hoechst as above. TMs were imaged on a Leica TCS SP8 confocal microscope. CFP, YFP, GFP, RFP, and Hoechst were detected using the following excitation/emission wavelengths, respectively: 458/463-490, 514/540-570, 488/490-510, 555/580-640, 405/410-500.

### TM explants and compound screen

For explant culture, murine TMs were floated in Advanced DMEM supplemented with 1% penicillin-streptomycin. They were maintained in an incubator at 37°C with 5% CO_2_. For drug exposures, the TMs were cultured in media with drug for 48 hours and then pulsed with 10 μM EdU for two hours. Whole-mounts were then processed as described above. Compounds tested in the screen are listed in **Table S3** (Selleckchem). Crenolanib used after the initial screen was from MedChem Express, and CP-673451 used after the initial screen was from Tocris.

### Human TM cell organoid cultures

Keratinocytes and fibroblasts were isolated from human TM tissue using procedures similar to ones previously described for isolation of cells from foreskin (Ridky et al. 2010). Briefly, the tissue was incubated in dispase at 4°C overnight, and then the epidermis and underlying tissue separated if possible. For keratinocyte isolation, the epidermis or bulk TM tissue was minced, then incubated in 0.5% trypsin (Life Technologies) at 37°C for 3 minutes. The enzyme was quenched with soybean trypsin inhibitor (Thermo Fisher Scientific), then the cells and tissue pelleted and subsequently plated. Keratinocytes were cultured in a 50/50 mix of Keratinocyte-SFM with supplements (EGF and BPE) and Medium 154 with HKGS (Thermo Fisher Scientific), plus 1% penicillin-streptomycin. For fibroblast isolation, the tissue was incubated in 1 mg/mL collagenase P in a 1:1:1 ratio of HBSS to dispase to fibroblast media (Advanced DMEM with 5% FBS and 1% penicillin-streptomycin) at 37°C for 1 hour. The tissue was then transferred to 0.5% trypsin and incubated at 37°C for 5 minutes. The digestion was quenched with fibroblast media, then both enzyme solutions were centrifuged and the cell pellets plated in fibroblast media.

The 3D organoid cultures were setup similarly to an assay previously described (Frank et al. 2016). We used KGM, a media previously employed for growth of organotypic skin cultures, which consists of a 3:1 mix of DMEM and Ham’s F12 supplemented with 10% FBS, 0.18 mM adenine (Sigma A9795), 0.4 μg/mL hydrocortisone (EMD Millipore 386698), 5 μg/mL insulin (Sigma I1882), 0.1 nM cholera toxin (EMD Millipore 227036), 10 ng/mL EGF (Life Technologies PHG0315), 0.01 mg/mL ciprofloxacin hydrochloride (Sigma PHR1044), 1.36 ng/mL triiodo-L-thyronine (Sigma T2877), 1% antibiotic-antimycotic, and 1% penicillin-streptomycin (Ridky et al. 2010). Per replicate, 8,000 cells (all keratinocytes, all fibroblasts, or a 50:50 mixture of the two) were resuspended in 45 μL KGM media, then 45 μL Matrigel (Corning 354234) was added and the resulting 90 μL plated in a transwell insert in a 24-well plate. The culture was allowed to set at 37°C for 20 minutes, then 600 μL of KGM media with 1 μM of the ROCK inhibitor thiazovivin (Selleckchem) was added to the well. After 24 hours, the media was changed to KGM without the ROCK inhibitor; for inhibitor studies, the compound was added at this point. Media was changed every 2-3 days. After 14 days, the cultures were fixed with 4% PFA at 4°C overnight, then washed with PBS. Images of the whole 3D cultures were captured on a Leica M205 FA microscope. The cultures were then equilibrated in 30% sucrose, embedded in OCT, and sectioned at 10 μm thickness for hematoxylin and eosin staining.

## DATA AVAILABILITY

All scRNA-Seq datasets have been deposited in NCBI’s Gene Expression Omnibus (GEO; accession number: GSE128892).

