## Supplemental Figures and Tables for "A hierarchy of migratory keratinocytes maintains the tympanic membrane"

Supplemental Figure Legends

**Figure S1, related to Figure 1: Proliferating cells captured in murine scRNA-Seq.** tSNE plots from the murine TM epidermis (top) and fibrous/mucosal fraction (bottom) showing expression of *Top2a* (left) and *Mki67* (right), two genes that mark cycling cells.

**Figure S2, related to Figure 1: *Fn1* expression in whole-mount TMs.** Left and middle: *in situ* hybridization using the RNAscope assay showing expression of *Fn1* (green), a marker undifferentiated keratinocyte cluster 2, in two whole-mount TMs. Right: A TM probed with negative control probes targeting bacterial transcripts is included on top.

**Figure S3, related to Figure 1: Visualization of cell types identified in the fibrous/mucosal scRNA-Seq dataset.** For each marker, expression in the fibrous/mucosal tSNE is shown on the left, and immunofluorescence in a whole-mount TM is shown on the right. Vim marks multiple mesenchymal clusters, Pecam1 marks endothelial cells (cluster 7), Acta2 marks smooth muscle cells (cluster 10), and Mbp marks Schwann cells (cluster 6).

**Figure S4, related to Figure 1: Expression of *Gpx3* and *Igfbp3* mark distinct mesenchymal cell populations.** (A) Heat-map showing expression of genes differentially expressed between cluster 0 and other mesenchymal cell clusters (2, 3, and 5). (B) Fibrous/mucosal fraction tSNE plots indicating expression of *Gpx3* and *Igfbp3*. (C) *In situ* hybridization indicating expression of *Gpx3* (cyan) and *Igfbp3* (green) in TM sections through the pars flaccida (top), upper pars tensa (middle), and mid pars tensa (bottom). Numbered boxes in the upper panels indicate the three areas magnified in the lower panels. In all sections, the epidermal layer is towards the top.

**Figure S5, related to Figure 1: Validation of mucosal cell markers from the scRNA-Seq dataset.** (A) Fibrous/mucosal fraction tSNE showing expression of *Sox2* (left), and immunofluorescence for Sox2 protein in a whole-mount TM (right). Gold boxes on the image of the entire whole-mount indicate the three areas magnified at the far right. (B) tSNE plots showing expression of *Dnah5* (left) and *Dynlrb2* (right). (C) *Krt19* expression in scRNA-Seq datasets from the fibrous/mucosal fraction (left) and epidermal fraction (middle). On the right, sections from a lineage traced Krt19-CreERT;mT/mG mouse.

**Figure S6, related to Figure 3: EdU pulse-chase demonstrates proliferative and migratory behaviors of TM cells.** The experimental time-line is indicated on top. On the bottom, whole-mount images of TMs harvested at the indicated time-points are shown. EdU is labeled in green.

**Figure S7, related to Figure 6: Lineage traced Krt5-CreERT2;R26R-Confetti TMs.** The TMs included in Figure 6 are shown with each channel (CFP, YFP, RFP, and GFP) broken out. Dense recombination was induced in Krt5-CreERT2;R26R-Confetti mice with 5 days of 2 mg/day tamoxifen, and then TMs were harvested and assessed at the time-points indicated to the left of the images.

**Figure S8, related to Figure 6: Quantification of color resolution in Krt5-CreERT2;R26R-Confetti TMs.** On top, and example of a TM harvested 3 months following tamoxifen administration is shown. To quantify the color mixing, a grid of 5,000 $\mu m^2$ (70.7 μm x 70.7 μm) boxes was superimposed over a maximum projection, and then the number of fluorescent proteins expressed in each box was manually annotated, as demonstrated in the magnified region of the TM. Very little GFP was seen, so a box could have 1, 2, or 3 colors. The results are shown graphically below.

<u>Supplemental Item Legends</u>

**Table S1, related to Figure 1. Gene sets from MSigDB enriched in cluster 1 relative to cluster 2.** The top 250 genes with increased expression in keratinocyte cluster 1 relative to keratinocyte cluster 2 were used to compute overlaps with gene sets in the Molecular Signatures Database. The top 50 gene sets are shown here.

**Table S2, related to Figure 1. Gene sets from MSigDB enriched in cluster 2 relative to cluster 1.** The top 250 genes with increased expression in keratinocyte cluster 2 relative to keratinocyte cluster 1 were used to compute overlaps with gene sets in the Molecular Signatures Database. The top 50 gene sets are shown here. The top 20 are also included in Figure 1M.

**Table S3, related to Figure 7. Compounds evaluated in the explant screen.** A custom library of small molecule inhibitors was purchased from Selleckchem. The inhibitor names and their protein targets are included in this table. This library was used to treat murine TMs in explant culture and assess for loss of proliferation. The results are in Figure 7B.

**Movie 1, related to Figure 4.** A Krt5-CreERT2;mT/mG mouse was given a single dose of 1 mg tamoxifen and a TM harvested two days later. It was maintained in Advanced DMEM in an incubator surrounding an AZ100 macro confocal microscope and imaged every hour for 24 hours. The resulting video loops four times. The first and last loops are not annotated, the second includes dragontails indicating where the cells have recently been, and the third includes the full tracks for each cell. Cells were tracked using the Spots function in Imaris.

**Movie 2, related to Figure 4.** A Ki67-CreERT2;mT/mG mouse was given a single dose of 250 mg/kg tamoxifen and a TM harvested two days later. It was maintained in Advanced DMEM in an incubator surrounding an AZ100 macro confocal microscope and imaged every hour for eight hours. The resulting video loops four times. The first and last loops are not annotated, the second includes dragontails indicating where the cells have recently been, and the third includes the full tracks for each cell. Cells were tracked using the Spots function in Imaris.

**Figure S1**

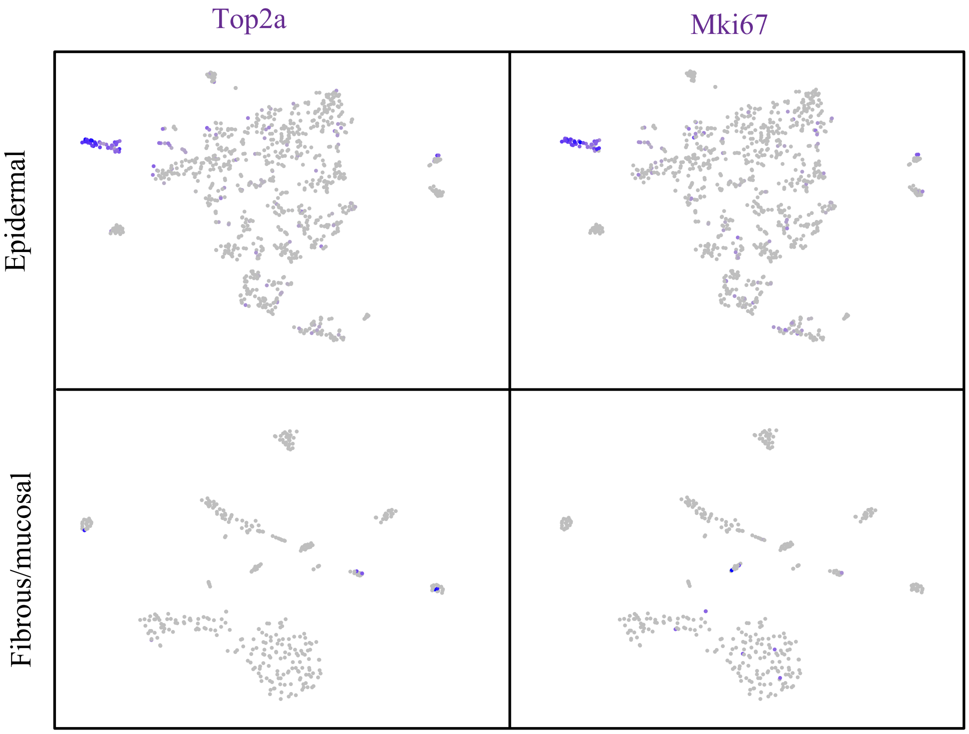

**Figure S2**

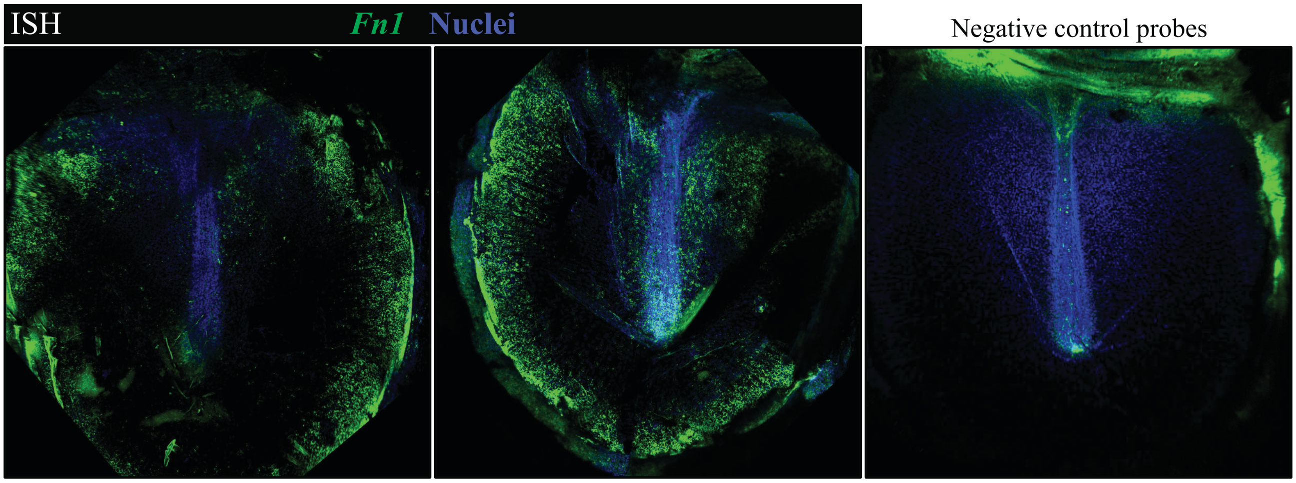

### Figure S3

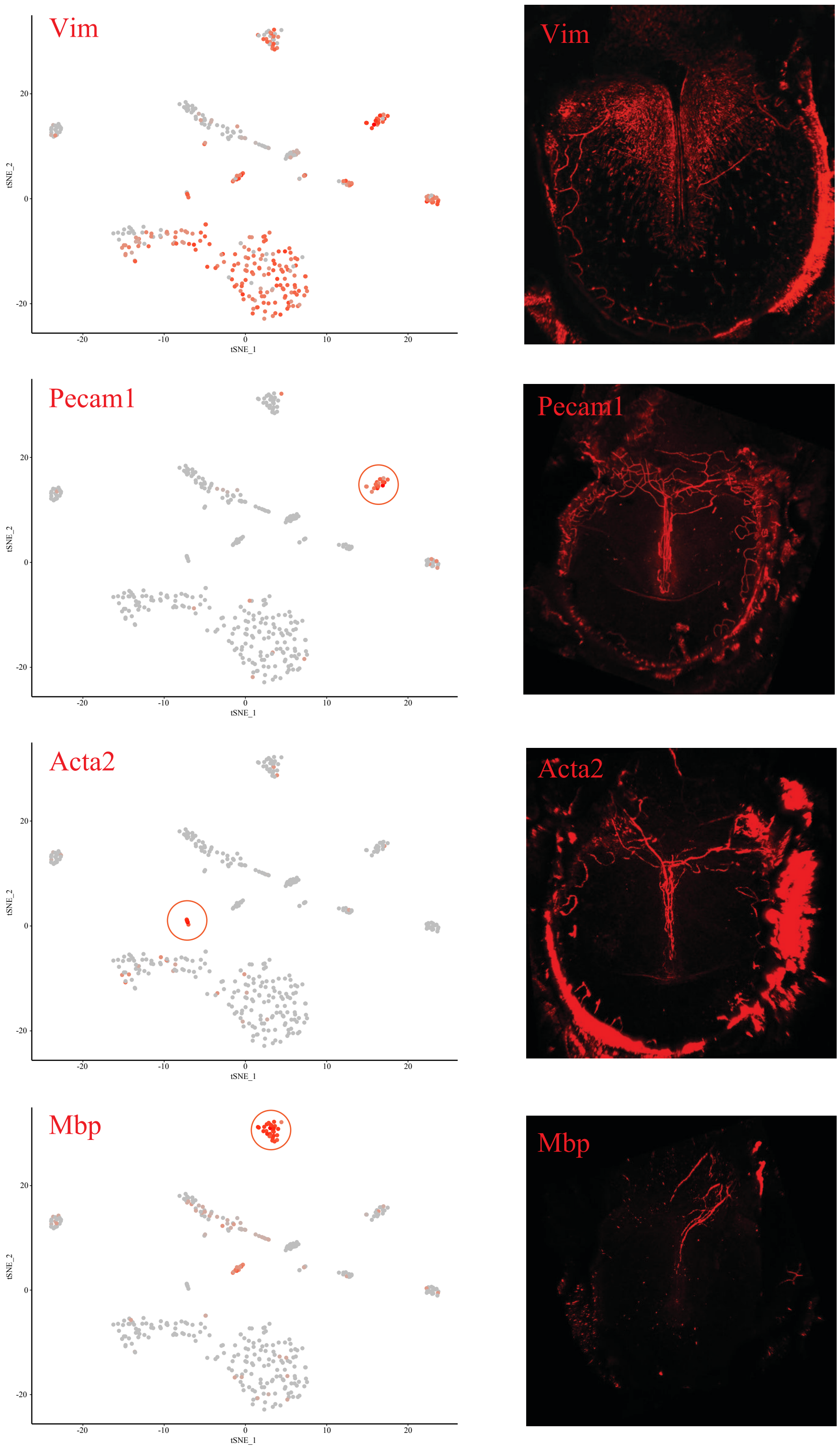
Vim
Vim
Pecam1
Pecam1
Acta2
Acta2
Mbp
Mbp
tSNE_1
tSNE_2
20
0
-20
-20
-10
0
10
20

### Figure S4

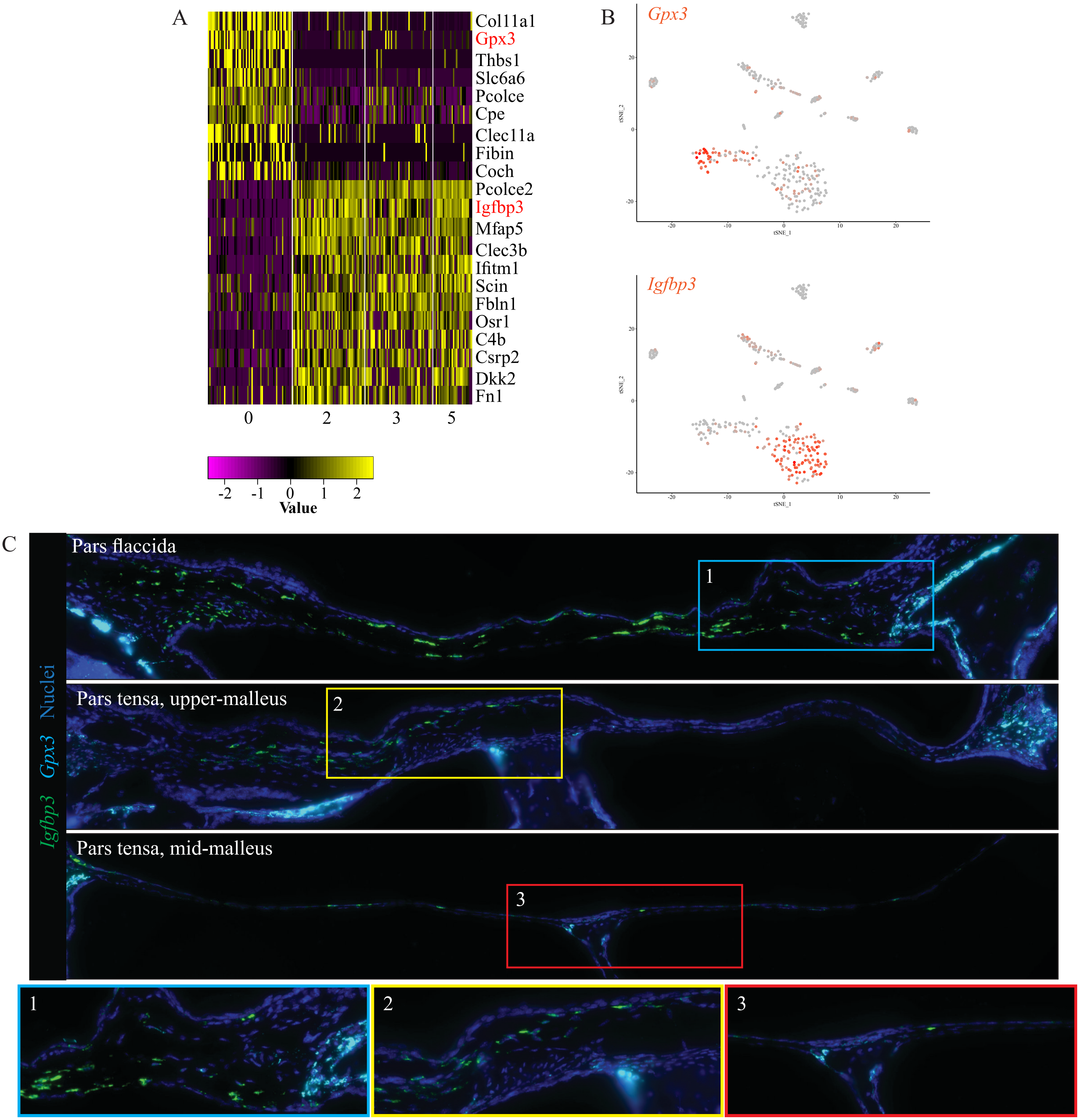
A
Col11a1
Gpx3
Thbs1
Slc6a6
Pcolce
Cpe
Clec11a
Fibin
Coch
Pcolce2
Igfbp3
Mfap5
Clec3b
Ifitm1
Scin
Fbln1
Osr1
C4b
Csrp2
Dkk2
Fn1
0
2
3
5
-2 -1 0 1 2
Value
B
Gpx3
Igfbp3
tSNE_1
tSNE_2
C
Igfbp3 Gpx3 Nuclei
Pars flaccida
Pars tensa, upper-malleus
Pars tensa, mid-malleus
1
2
3

**Figure S5**

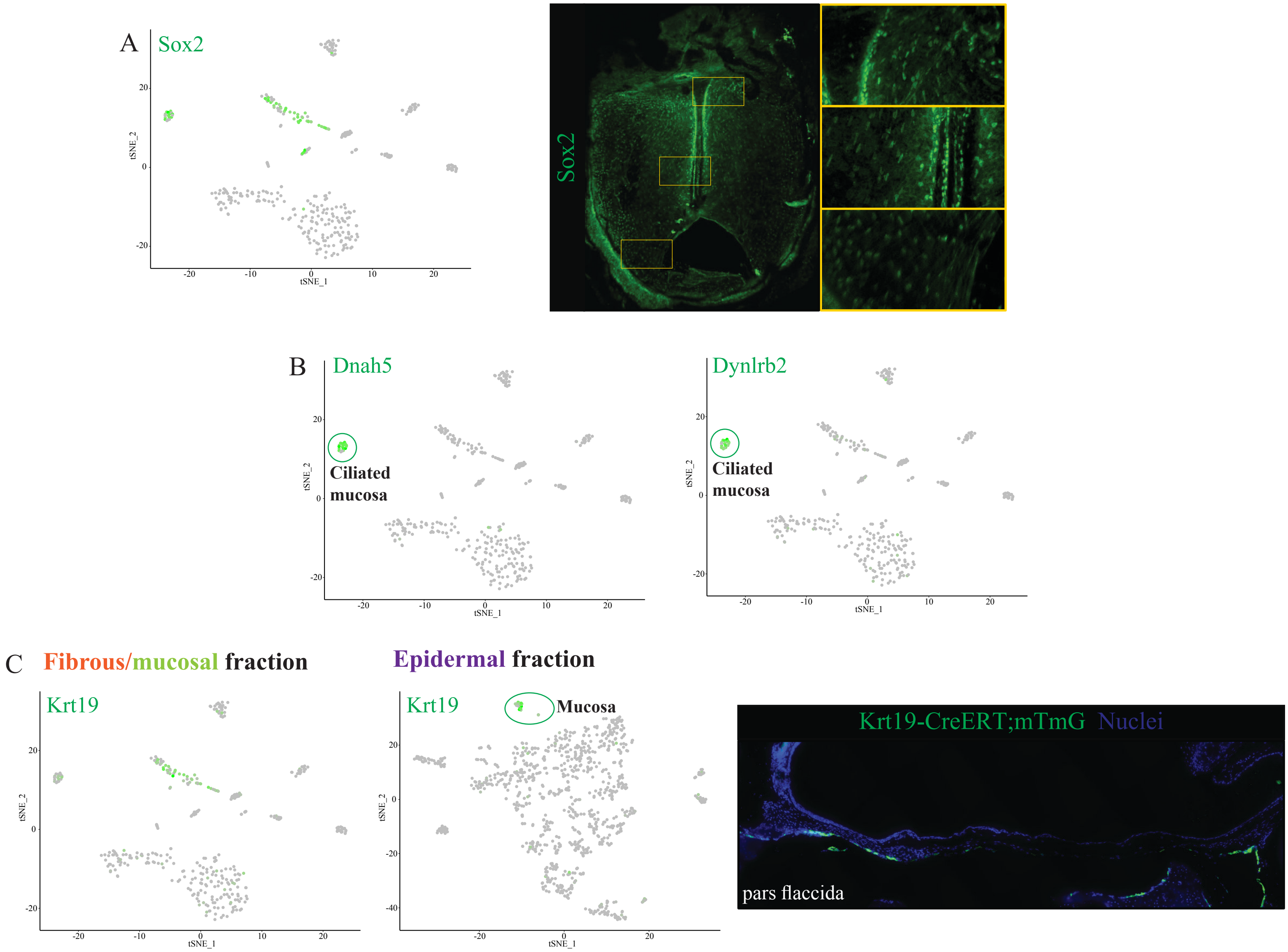

A
Sox2
tSNE_2
tSNE_1
Sox2
B
Dnah5
Ciliated mucosa
Dynlrb2
Ciliated mucosa
C
Fibrous/mucosal fraction
Epidermal fraction
Krt19
Krt19
Mucosa
Krt19-CreERT;mTmG Nuclei
pars flaccida

**Figure S6**

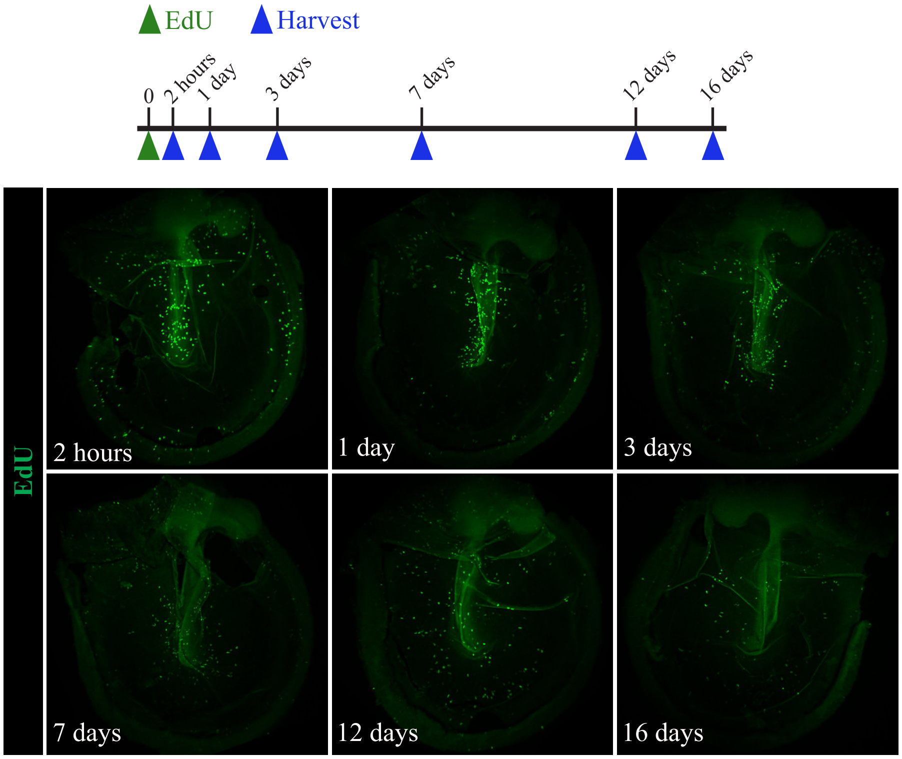
EdU
Harvest
0
2 hours
1 day
3 days
7 days
12 days
16 days
EdU
2 hours
1 day
3 days
7 days
12 days
16 days

**Figure S7**

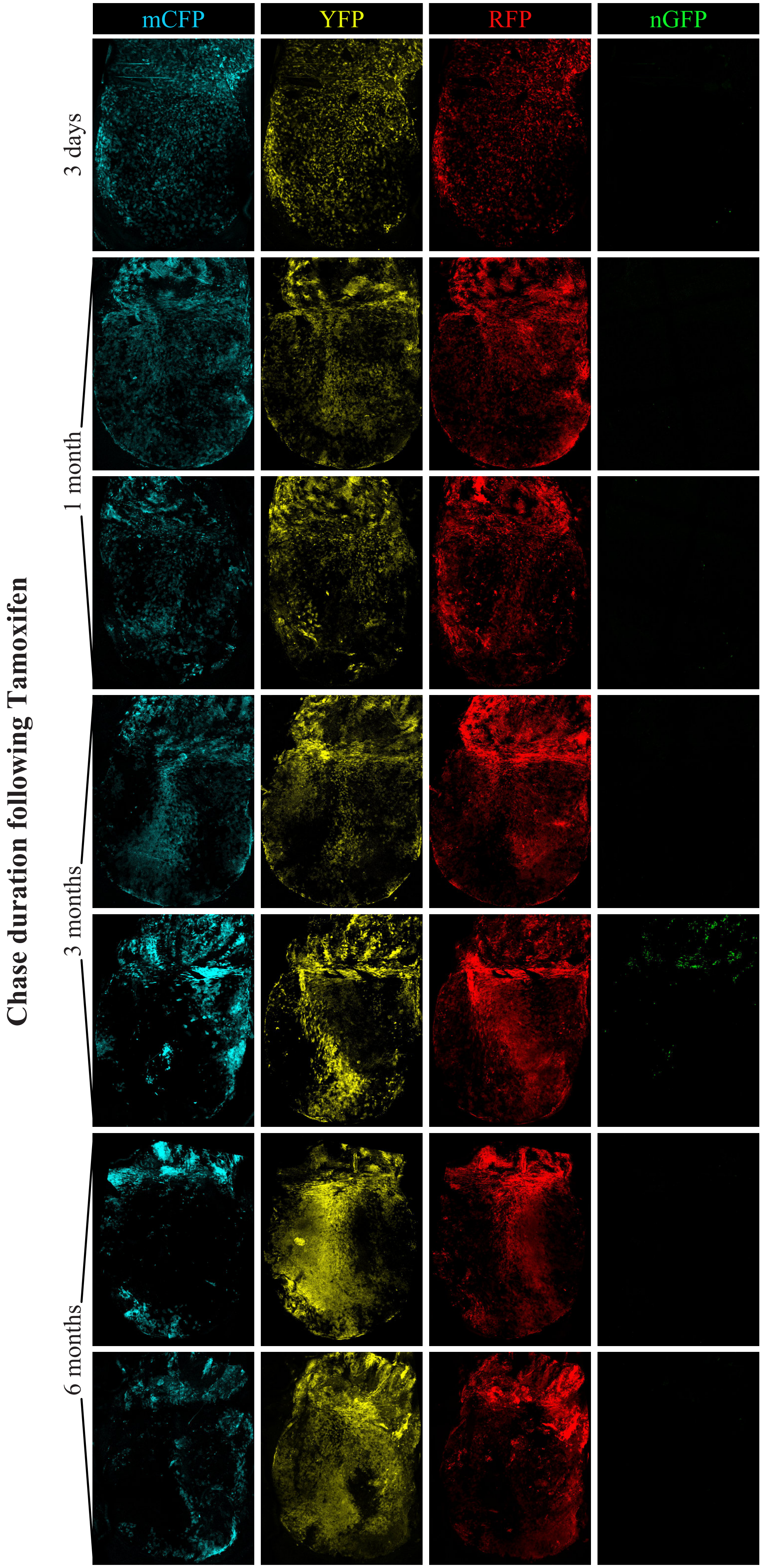
mCFP
YFP
RFP
nGFP
3 days
1 month
3 months
6 months
Chase duration following Tamoxifen

**Figure S8**

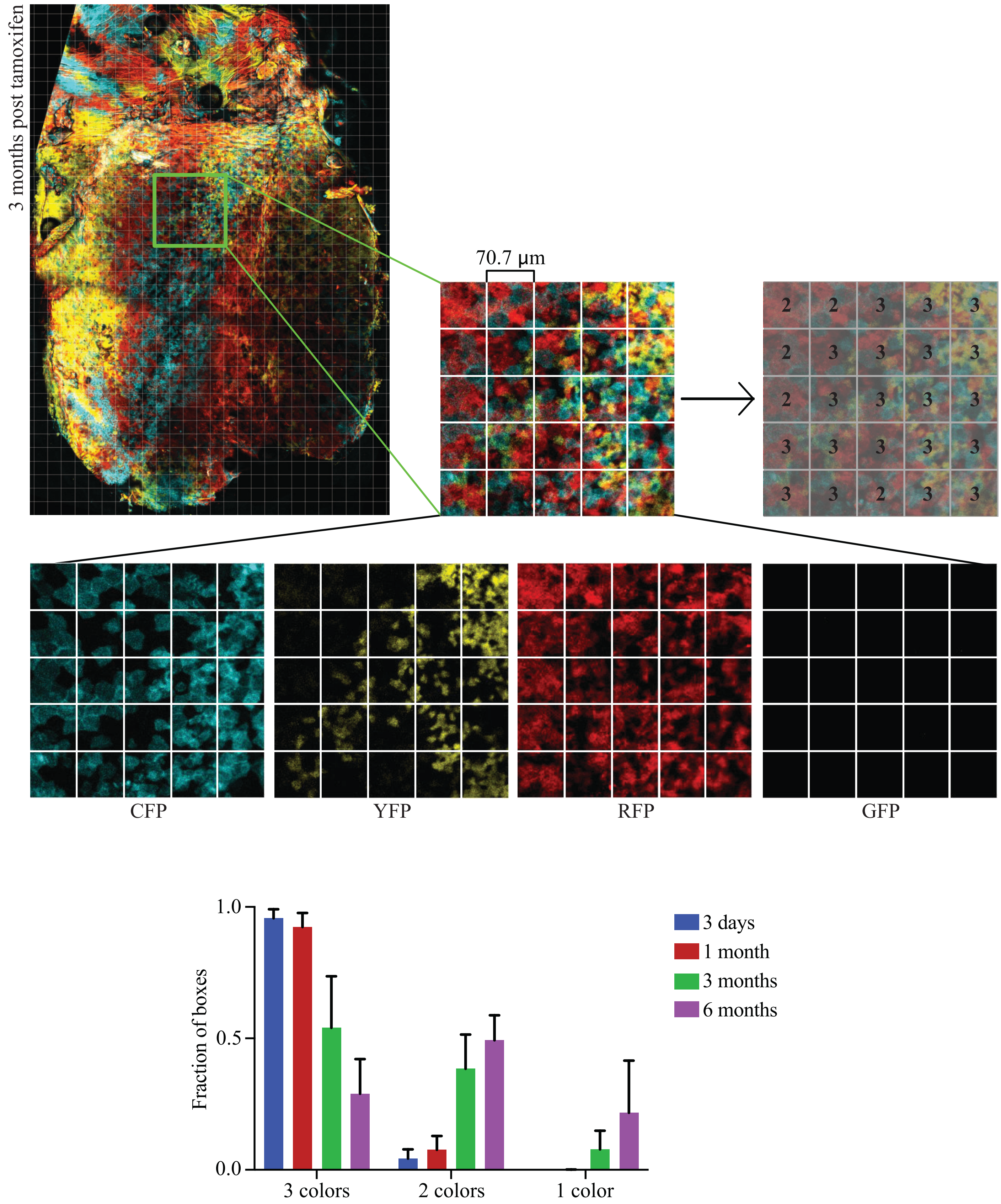
3 months post tamoxifen
70.7 µm
2 2 3 3 3
2 3 3 3 3
2 3 3 3 3
3 3 3 3 3
3 3 2 3 3
CFP
YFP
RFP
GFP
Fraction of boxes
1.0
0.5
0.0
3 colors
2 colors
1 color
3 days
1 month
3 months
6 months

**Table S1, related to Figure 1: Gene sets from MSigDB enriched in cluster 1 relative to cluster 2.**

| Gene Set Name | p-value |
|---|---|
| KEGG_RIBOSOME | 7.49E-153 |
| GO_ESTABLISHMENT_OF_PROTEIN_LOCALIZATION_TO_ENDOPLASMIC_RETICULUM | 1.62E-143 |
| GO_CYTOSOLIC_RIBOSOME | 4.05E-142 |
| GO_NUCLEAR_TRANSCRIBED_MRNA_CATABOLIC_PROCESS_NONSENSE_MEDIATED_DECAY | 5.09E-140 |
| GO_TRANSLATIONAL_INITIATION | 6.47E-138 |
| GO_PROTEIN_LOCALIZATION_TO_ENDOPLASMIC_RETICULUM | 1.93E-135 |
| GO_MULTI_ORGANISM_METABOLIC_PROCESS | 7.12E-133 |
| GO_RIBOSOMAL_SUBUNIT | 3.14E-128 |
| REACTOME_3_UTR_MEDIATED_TRANSLATIONAL_REGULATION | 2.03E-127 |
| GO_PROTEIN_TARGETING_TO_MEMBRANE | 2.19E-127 |
| REACTOME_PEPTIDE_CHAIN_ELONGATION | 4.90E-126 |
| REACTOME_TRANSLATION | 9.03E-125 |
| REACTOME_INFLUENZA_VIRAL_RNA_TRANSCRIPTION_AND_REPLICATION | 5.00E-122 |
| REACTOME_NONSENSE_MEDIATED_DECAY_ENHANCED_BY_THE_EXON_JUNCTION_COMPLEX | 1.97E-120 |
| REACTOME_SRP_DEPENDENT_COTRANSLATIONAL_PROTEIN_TARGETING_TO_MEMBRANE | 9.00E-120 |
| REACTOME_INFLUENZA_LIFE_CYCLE | 3.48E-117 |
| GO_RIBOSOME | 2.69E-115 |
| HSIAO_HOUSEKEEPING_GENES | 9.85E-115 |
| GO_CYTOSOLIC_PART | 1.23E-113 |
| GO_STRUCTURAL_CONSTITUENT_OF_RIBOSOME | 2.39E-113 |
| GO_RNA_CATABOLIC_PROCESS | 5.67E-113 |
| GO_RRNA_METABOLIC_PROCESS | 7.05E-113 |
| GO_ESTABLISHMENT_OF_PROTEIN_LOCALIZATION_TO_MEMBRANE | 1.40E-111 |
| GO_RIBOSOME_BIOGENESIS | 6.47E-110 |
| REACTOME_METABOLISM_OF_MRNA | 7.54E-107 |

| Gene Set Name | p-value |
|---|---|
| GO_VIRAL_LIFE_CYCLE | 4.36E-104 |
| GO_PROTEIN_LOCALIZATION_TO_MEMBRANE | 1.44E-102 |
| GO_AMIDE_BIOSYNTHETIC_PROCESS | 1.59E-102 |
| GO_RIBONUCLEOPROTEIN_COMPLEX_BIOGENESIS | 2.11E-102 |
| REACTOME_METABOLISM_OF_RNA | 1.74E-101 |
| GO_ESTABLISHMENT_OF_PROTEIN_LOCALIZATION_TO_ORGANELLE | 3.13E-100 |
| GO_PROTEIN_TARGETING | 8.53E-100 |
| GO_PEPTIDE_METABOLIC_PROCESS | 9.28E-100 |
| GO_NCRNA_PROCESSING | 9.73E-100 |
| BILANGES_SERUM_AND_RAPAMYCIN_SENSITIVE_GENES | 2.20E-97 |
| REACTOME_METABOLISM_OF_PROTEINS | 2.15E-96 |
| GO_ORGANIC_CYCLIC_COMPOUND_CATABOLIC_PROCESS | 2.38E-94 |
| GO_CELLULAR_AMIDE_METABOLIC_PROCESS | 6.52E-94 |
| GO_INTERSPECIES_INTERACTION_BETWEEN_ORGANISMS | 7.74E-91 |
| GO_ORGANONITROGEN_COMPOUND_BIOSYNTHETIC_PROCESS | 5.56E-90 |
| GO_PROTEIN_LOCALIZATION_TO_ORGANELLE | 1.06E-88 |
| GO_NCRNA_METABOLIC_PROCESS | 1.84E-88 |
| GO_CYTOSOLIC_LARGE_RIBOSOMAL_SUBUNIT | 4.15E-87 |
| GO_RIBONUCLEOPROTEIN_COMPLEX | 1.59E-84 |
| GO_INTRACELLULAR_PROTEIN_TRANSPORT | 2.88E-83 |
| GO_STRUCTURAL_MOLECULE_ACTIVITY | 2.76E-79 |
| GO_MRNA_METABOLIC_PROCESS | 6.08E-79 |
| GO_SINGLE_ORGANISM_CELLULAR_LOCALIZATION | 2.49E-78 |
| GO_MACROMOLECULE_CATABOLIC_PROCESS | 2.98E-77 |
| GO_ORGANONITROGEN_COMPOUND_METABOLIC_PROCESS | 5.90E-77 |

**Table S2, related to Figure 1: Gene sets from MSigDB enriched in cluster 2 relative to cluster 1.**

| Gene Set Name | p-value | Gene Set Name | p-value |
|---|---|---|---|
| WONG_ADULT_TISSUE_STEM_MODULE | 1.21E-40 | GO_REGULATION_OF_MULTICELLULAR_ORGANISMAL_DEVELOPMENT | 1.01E-25 |
| MEISSNER_BRAIN_HCP_WITH_H3K4ME3_AND_H3K27ME3 | 1.81E-40 | MCBRYAN_PUBERTAL_BREAST_4_5WK_UP | 1.28E-25 |
| GO_TISSUE_DEVELOPMENT | 1.17E-38 | GO_NEUROGENESIS | 2.12E-25 |
| GO_EXTRACELLULAR_MATRIX | 6.27E-36 | GO_REGULATION_OF_CELL_PROLIFERATION | 3.98E-25 |
| GO_EXTRACELLULAR_STRUCTURE_ORGANIZATION | 1.45E-35 | LIU_PROSTATE_CANCER_DN | 4.22E-25 |
| MILI_PSEUDOPODIA_HAPTOTAXIS_DN | 1.22E-34 | LEE_BMP2_TARGETS_UP | 9.34E-25 |
| PASINI_SUZ12_TARGETS_DN | 1.40E-33 | JOHNSTONE_PARVB_TARGETS_3_UP | 4.36E-24 |
| REN_ALVEOLAR_RHABDOMYOSARCOMA_DN | 3.45E-31 | NABA_CORE_MATRISOME | 4.37E-24 |
| GO_BIOLOGICAL_ADHESION | 3.85E-31 | WU_CELL_MIGRATION | 5.69E-24 |
| MARTINEZ_RB1_AND_TP53_TARGETS_DN | 1.31E-30 | GO_TISSUE_MORPHOGENESIS | 9.02E-24 |
| GO_CELL_SURFACE | 4.53E-30 | BLALOCK_ALZHEIMERS_DISEASE_UP | 9.81E-24 |
| GO_EXTRACELLULAR_SPACE | 1.31E-29 | JAEGER_METASTASIS_DN | 2.11E-23 |
| GO_ORGAN_MORPHOGENESIS | 2.13E-29 | MILI_PSEUDOPODIA_CHEMOTAXIS_DN | 2.41E-23 |
| GO_MOVEMENT_OF_CELL_OR_SUBCELLULAR_COMPONENT | 3.92E-29 | GO_EPITHELIUM_DEVELOPMENT | 2.73E-23 |
| GO_LOCOMOTION | 1.19E-28 | HALLMARK_EPITHELIAL_MESENCHYMAL_TRANSITION | 3.71E-23 |
| LIM_MAMMARY_STEM_CELL_UP | 1.48E-28 | GO_CELL_JUNCTION | 3.88E-23 |
| GO_REGULATION_OF_HYDROLASE_ACTIVITY | 2.35E-28 | GO_POSITIVE_REGULATION_OF_CATALYTIC_ACTIVITY | 4.65E-23 |
| GO_RESPONSE_TO_OXYGEN_CONTAINING_COMPOUND | 1.39E-27 | BAELDE_DIABETIC_NEPHROPATHY_DN | 9.19E-23 |
| NABA_MATRISOME | 5.30E-27 | GO_POSITIVE_REGULATION_OF_MOLECULAR_FUNCTION | 1.15E-22 |
| MARTINEZ_TP53_TARGETS_DN | 5.72E-27 | GO_REGULATION_OF_CELLULAR_COMPONENT_MOVEMENT | 3.75E-22 |
| GO_RESPONSE_TO_ENDOGENOUS_STIMULUS | 1.21E-26 | GO_CELLULAR_RESPONSE_TO_ORGANIC_SUBSTANCE | 4.33E-22 |
| GO_PROTEINACEOUS_EXTRACELLULAR_MATRIX | 2.03E-26 | GO_CELLULAR_COMPONENT_MORPHOGENESIS | 5.15E-22 |
| GO_INTRACELLULAR_VESICLE | 2.07E-26 | GO_CELL_MORPHOGENESIS_INVOLVED_IN_DIFFERENTIATION | 6.01E-22 |
| GO_REGULATION_OF_CELL_DIFFERENTIATION | 4.24E-26 | GO_NEURON_DIFFERENTIATION | 1.96E-21 |
| GO_CELL_DEVELOPMENT | 5.07E-26 | GO_ANCHORING_JUNCTION | 2.28E-21 |

**Table S3, related to Figure 7: Compounds evaluated in the explant screen.**

| Product Name | Target(s) |
|---|---|
| Risperidone | 5-HT Receptor |
| BRL-15572(dihydrochloride) | 5-HT Receptor |
| Istradefylline | Adenosine Receptor |
| SCH58261 | Adenosine Receptor |
| Timolol Maleate | Adrenergic Receptor |
| ICI-118551 Hydrochloride | Adrenergic Receptor |
| A-769662 | AMPK |
| Tozasertib (VX-680, MK-0457) | Aurora Kinase |
| PD173955 | Bcr-Abl |
| GNF-5 | Bcr-Abl |
| RO4929097 | Beta Amyloid,Gamma-secretase |
| MK-0752 | Beta Amyloid,Gamma-secretase |
| PHA-665752 | c-Met |
| SU11274 | c-Met |
| Palbociclib (PD-0332991) HCl | CDK |
| SNS-032 (BMS-387032) | CDK |
| AG-1478 (Tyrphostin AG-1478) | EGFR |
| PD168393 | EGFR |
| CP-724714 | EGFR,HER2 |
| Bosentan Hydrate | Endothelin Receptor |
| Sitaxentan sodium | Endothelin Receptor |
| Zibotentan (ZD4054) | Endothelin Receptor |
| Fulvestrant | Estrogen/progestogen Receptor |
| PF-573228 | FAK |
| TAE226 (NVP-TAE226) | FAK |
| AZD4547 | FGFR |
| SSR128129E | FGFR |
| LY411575 | Gamma-secretase |
| Semagacestat (LY450139) | Gamma-secretase |
| CHIR-99021 (CT99021) | GSK-3 |
| SB216763 | GSK-3 |
| SB415286 | GSK-3 |
| TWS119 | GSK-3 |
| BMS-833923 | Hedgehog/Smoothened |
| PF-5274857 | Hedgehog/Smoothened |
| GSK1904529A | IGF-1R |
| Linsitinib (OSI-906) | IGF-1R |
| IMD 0354 | IκB/IKK |
| TPCA-1 | IκB/IKK |
| Decernotinib (VX-509) | JAK |
| Ruxolitinib (INCB018424) | JAK |
| (+)-MK 801 maleate | NMDAR |
| FRAX597 | PAK |
| CP-673451 | PDGFR |
| Crenolanib (CP-868596) | PDGFR |
| Enzastaurin (LY317615) | PKC |
| Go 6983 | PKC |
| GW9662 | PPAR |
| FH535 | PPAR,Wnt/beta-catenin |
| Fasudil (HA-1077) HCl | ROCK |
| GSK429286A | ROCK |
| Thiazovivin | ROCK |
| Y-27632 2HCl | ROCK |
| Ozanimod (RPC1063) | S1P Receptor |
| PF-543 | S1P Receptor |
| Bosutinib (SKI-606) | Src |
| APTSTAT3-9R | STAT |
| HO-3867 | STAT |
| SH-4-54 | STAT |
| R406 (free base) | Syk |
| LDN-193189 HCl | TGF-beta/Smad |
| RepSox | TGF-beta/Smad |
| SB431542 | TGF-beta/Smad |
| K02288 | TGF-beta/Smad |
| GSK2193874 | Trpv4 inhibitor |
| SAR131675 | VEGFR |
| Lenvatinib (E7080) | VEGFR |
| ICG-001 | Wnt/beta-catenin |
| IWP-L6 | Wnt/beta-catenin |
| LGK-974 | Wnt/beta-catenin |
| PRI-724 | Wnt/beta-catenin |
| Wnt agonist 1 | Wnt/beta-catenin |
| Wnt-C59 (C59) | Wnt/beta-catenin |
| XAV-939 | Wnt/beta-catenin |
